# Membrane-coupled conformational switching of VISTA regulates immune checkpoint signaling via CC′ loop accessibility

**DOI:** 10.64898/2026.02.07.704574

**Authors:** Aravindhan Ganesan, Varun Reddy, Norman Ly

## Abstract

V-domain Ig Suppressor of T cell activation (VISTA) has emerged as a critical target for anti-cancer immunotherapy. VISTA inhibits T cell activity, suppressing the anti-cancer immune responses. Here, we use atomistic molecular dynamics simulations to investigate the conformational behavior of membrane-bound human VISTA using both a glycan-free truncated model and a glycosylated, full-length model. Our simulations show that the extracellular and transmembrane domains remain structurally stable; notably, the unusually long CC′ loop is stabilized by extensive hydrogen-bonding interactions. Principal component analysis and conformational clustering reveal a dominant rotational motion of the extracellular domain relative to the membrane, giving rise to two recurrent conformational states: an “Up” state in which the CC′ loop is solvent-exposed and accessible for ligand engagement, and a “Down” state in which the loop transiently associates with the lipid bilayer. These transitions are allosterically coupled to proline-mediated bending of the transmembrane helix. We propose that this membrane-coupled Up/Down conformational switching regulates CC′ loop accessibility, favoring cis interactions with cognate protein partners on the same cell surface, consistent with recent experimental observations, while permitting context-dependent trans interactions. Together, our findings reveal a membrane-coupled conformational mechanism regulating VISTA immune checkpoint function.

## 1. Introduction

The advent of immune checkpoints has revolutionized the field of cancer immunotherapy.^1^ According to the two-signal model of T-cell activation, the primary signal is generated by the T-cell receptor’s recognition of a specific peptide presented by the Major Histocompatibility Complex (MHC) on an antigen-presenting cell (APC). A secondary signal, delivered through specific interactions between integral immunoglobulin (Ig) superfamily proteins, determines the fate of T-cell activation. For instance, the binding of CD28 on T cells with B7 ligands (B7-1 and B7-2) on APCs delivers a co-stimulatory signal that promotes T-cell activation and proliferation. In contrast, other Ig proteins, generally known as immune checkpoints, such as cytotoxic T-lymphocyte-associated antigen (CTLA-4) and programmed cell death protein 1 (PD-1), with their complementary ligands (B7-1/B7-2 for CTLA-4 and PD-L1/PD-L2 for PD-1), induce T-cell inactivation to maintain homeostasis and self-tolerance.^2, 3^ Cancers exploit these checkpoints by overexpressing them to evade immune surveillance and grow unchecked.^4-6^ Monoclonal antibodies (mAbs) targeting these specific immune checkpoints effectively block their signaling to re-activate T-cell activity for tumor clearance.^7^ Therefore, the discovery and structural characterization of novel checkpoint proteins and the development of targeted blockade strategies (mAbs and small molecules) have become a crucial area of research.

V-domain lg suppressor of T cell activation (VISTA) is a novel inhibitory immune checkpoint protein overexpressed in various cancers, including prostate and breast cancers. Encoded by Cdh23 gene on chromosome 10 (10q22.1), VISTA is highly expressed in the myeloid cells (e.g., dendritic cells and neutrophils) and promotes the inhibition of T cell activity and the downstream inflammatory cytokines such as interleukin-2 and interferon-γ required for anti-cancer immune responses.^4^ VISTA has also been shown to be upregulated in some patients treated with mAbs targeting other checkpoints (such as CTLA-4 and PD-L1), promoting non-responsiveness and resistance to current therapeutic modalities.^8, 9^ As a result, VISTA has emerged as an important therapeutic target against cancers.

VISTA is an integral membrane protein belonging to the B7:CD28 family. It is a 279-amino acid protein (excluding a 32-residue signal peptide) composed of three domains: a 162-residue extracellular domain (ECD), a 21-residue transmembrane domain (TMD), and a 96-residue intracellular domain (ICD). VISTA displays sequence-structure-function uniqueness when compared to other known Ig checkpoints. Unlike the other known checkpoints, VISTA can function as both a receptor on the surface of T cells or a ligand on the APC surface.^10^ Until date, four experimentally resolved X-ray crystal structures of VISTA’s ECD in both apo-states (Protein data bank or PDB codes: 6U6V^11^; 6OIL^12^) and mAB-bound complexes (PDB codes: 6MVL^13^; 8TBQ^14^) have been reported in the PDB, Figure 1A. These structures reveal that VISTA’s ECD has a single large Ig-V domain preserving an anti-parallel β-sheets arrangement that is composed of 10 β-sheets (named as A-H in Figure 1B) interconnected by loops and 3 helices. Similar to other checkpoints, VISTA’s ECD has two faces, namely, the receptor binding face (A’BEDC”) on one side and the ligand binding face (HAGFCC’) on the opposite side.^10^ Nevertheless, the presence of an extra 10^th^ strand (the H strand) in VISTA makes it unique when compared to the other checkpoints. The last H strand in VISTA ECD is connected to its TMD through a stalk segment.^10^ While other checkpoints have only a single disulfide bond in their ECDs, the structure of VISTA ECD is stabilized by three disulfide bonds formed by C12^A-strand^: C146^H-strand^, C51^CC’loop^:C113^F-strand^, and C22^B-strand^:C114^F-strand^ pairs (Figure 1C). Two of these pairs (C12^A-strand^: C146^H-strand^, C51^CC’loop^:C113^F-strand^) are unique to VISTA and these covalent linkages connect the A and H strands, as well as the CC’ loop to the F strand.^10^ Our recent molecular dynamics (MD) simulation study^1^ on the VISTA: mAb complexes revealed key arginine fingerprint residues on CC’ loop and FG loop played a critical role in its interactions with pH-selective mAbs. In vitro mutagenesis of these disulfide bonds and deletion of the unique H strand in VISTA ECD resulted in the loss of inhibitory activity.^1^

**Figure 1.**
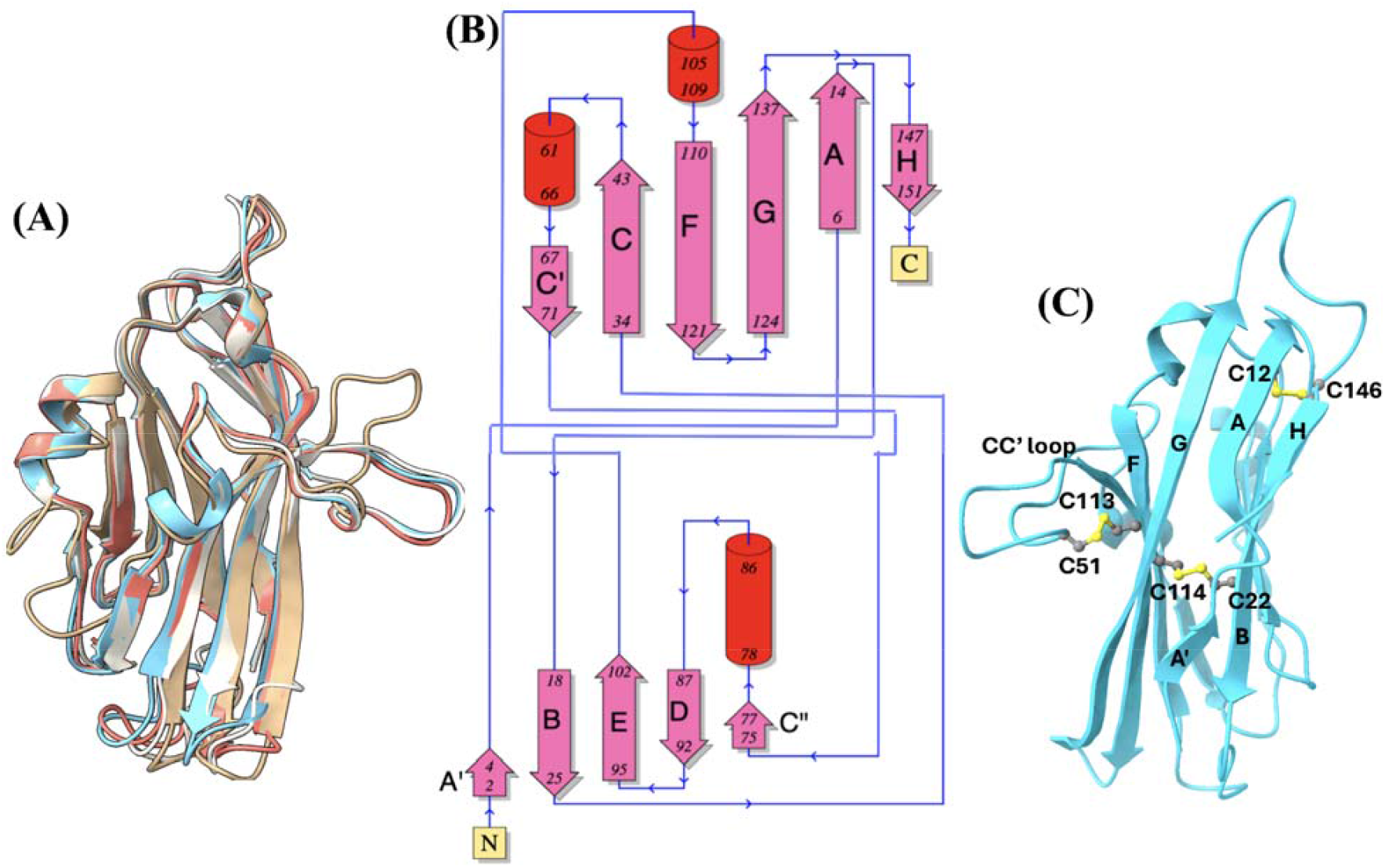
3D structures and 2D topology diagram of human VISTA ECD. (A) Superimposed view of all the four VISTA ECD X-ray crystal structures reported in the PDB (PDBs: 6U6V, 6OIL, 6MVL, 8TBQ) shown in a cartoon representation. The missing loops in the crystal structures were modelled using Modeller program available as plugin within the ChimeraX program, and the 6U6V structure was used as reference fit the other structures. The overlapped image shows high levels of similarity in their orientations, except for a few displacements in the loops. (B) The 2D structural topology diagram of VISTA ECD generated using the PDBSum program and slightly modified for visual purpose. The 2D diagram describes the labels of the beta sheets (in pink), along with the helices (red) and interconnecting loops (in blue). (C) 3D structure of VISTA ECD (blue cartoon) showing the three disulfide bonds (ball and sticks) formed by C12^A-strand^: C146^H-strand^, C51^CC’loop^:C113^F-strand^, and C22^B-strand^:C114^F-strand^. The cysteine residues and their corresponding secondary structural elements are labelled in the image.

Another unique feature to VISTA among other checkpoints is the presence of surprisingly large amount of histidine (HIS) residues in its ECD. VISTA ECD is made up of 8.6% of HIS residues giving it an ability to bind different binding partners under distinct pH conditions. It has been shown that VISTA ECD is able to interact with V-Set and Immunoglobin domain containing-3 (VSIG-3) expressed on APCs in neutral pH^14-17^, and with P-selecting glycoprotein ligand-1 (PSGL-1) expressed on the T cell surface in acidic pH conditions.^18-19^ Interactions of VISTA with its binding partners at the immunological synapse has been shown to inhibit T cell function, hindering cytokine production. Genetic knockout of VISTA in a mouse model resulted in the accumulation of activated T-cells and, consequently, the generation of various inflammatory cytokines and chemokines.^20^ Thus, VISTA plays a critical role in the regulation of T cell mediated immune responses and is a promising target for cancer immunotherapy.

While the structure of VISTA ECD has been experimentally reported, the full model of VISTA is only accessible through the AI-predictions in AlphaFold^21^. While the TMD of VISTA in this AlphaFold model (AF-Q9H7M9-F1-v4) is predicted with moderate confidence, its ICD is modelled with very low confidence and as a highly disordered loop that is misoriented towards the TMD of VISTA (Supplementary Information, Figure S1). As a result, there is a need for a high-quality, comprehensive model of VISTA to understand its conformational changes in solution. Such insights will be useful to expand our understanding on the structure-dynamics of VISTA relevant for its signaling processes.

To address this gap, this work focuses on building a comprehensive model of human VISTA using a combination of molecular modelling approaches and optimizing the model in a membrane-embedded environment through extensive all-atom MD simulations. Our MD analyses reveal that VISTA exhibits an interesting conformational switching between two states, dubbed as ‘Up’ and ‘Down’ states, mediated through its interactions with the lipid bilayer. In the Up state, VISTA is oriented away from the membrane such that its ligand-binding face is exposed; while in the Down state, VISTA ECD bends towards the TMD, allowing its interactions with the lipid bilayer. To confirm that the conformational state is not biased by the starting conformation or the MD program employed, the full model of glycosylated VISTA (without any truncation) was modelled and simulated independently using a different MD engine. This complete model also sampled these distinct ECD rotations, suggesting that this conformational switching is physiologically relevant for VISTA. Our findings provide novel mechanistic insights into VISTA, suggesting a lipid-mediated regulatory mechanism for its ligand binding.

## 2. Methodology

This work employs a multi-step approach, combining AI-based structure prediction (AlphaFold)^21^ with ab initio modeling and membrane building to construct a comprehensive atomistic model of the human VISTA protein integrated into a lipid bilayer. The final model was optimized with three independent 500 ns long classical MD simulations, and the conformational dynamics of VISTA were examined as described below.

### 2.1. Atomistic model building of VISTA

While the three-dimensional (3D) X-ray crystal structures of the VISTA ECD are available, the full-length VISTA model was accessed from the AlphaFold database, predicted through an AI approach^21^. The AlphaFold model (AF-Q9H7M9-F1-v4) is shown in the Supplementary Information, Figure S1. Although the ECD and TMD segments were predicted with high to very high confidence, the ICD had a poor resolution with a very low model confidence score. Furthermore, the ICD was completely disordered and misoriented toward the TMD, which presented challenges for membrane embedding. To address this, Rosetta ab initio modeling^22^ was used to generate a 3D structure of the VISTA ICD. However, the ab initio models of the full-length VISTA ICD (96 residues spanning Y184-I279) also displayed a poor model confidence score (0.19), which is not surprising given the disordered nature of the ICD. Therefore, the last 51 residues of the ICD were omitted, and only the first 45 residues (spanning Y184-R228) were remodeled using Rosetta, which slightly improved the confidence score to 0.25. This truncated ICD model was then used to replace the full ICD in the AlphaFold structure to build a comprehensive final VISTA model (shown in Supplementary Information, Figure S2). The final model (herein referred to as the VISTA model in the text) was used for probing its conformational dynamics using MD simulation. Because the first 32 residues corresponding to the signal peptide were excluded from the structural model, the numbering in this study is renormalized such that residue 33 of the full-length sequence is referred to as residue 1.

### 2.2. Classical MD simulation of membrane-bound VISTA

#### System preparation

The protonation states of ionizable residues in the final model were assigned at a neutral pH of 7.4 using the PROPKA 3.1 PDB2PQR server^23, 24^. The prepared protein was subsequently embedded into a lipid bilayer composed of 1-Palmitoyl-2-oleoyl-sn-glycero-3-phosphocholine (POPC), a common zwitterionic phospholipid widely used as a model for biological membranes.^25^ The membrane-integrated VISTA model was solvated in an aqueous box with a 22.5 Å buffer distance from the protein to the box edges, using TIP3P water molecules. The system was then electro neutralized and a salt concentration of 0.15 M KCl was added. All steps of membrane integration and solvation were carried out using the CHARMM-GUI program.^26, 27^

#### Simulation protocol

The prepared VISTA model was subjected to classical MD simulation using the NAMD 3.0.1 program. The VISTA structure was described using the CHARMM36m protein force field, the POPC lipids were describe using the CHARMM36 lipid force field parameters. Initially, the model system was energy minimized in 30,000 steps, followed by a simulation with an NVT ensemble (constant Number of particles, Volume, and Temperature) to heat the system to 310 K, using a Langevin dynamics over 2 ns. This was followed by 6 steps of equilibration with the NPT ensemble (constant Number of particles, Pressure and Temperature) at 310 K and 1 bar using the Nose-Hoover thermostat, and the Parrinello-Rahman barostat.^28^ Harmonic restraints on the protein’s backbone and sidechain atoms were gradually reduced over six steps: 10 kcal mol□^1^ Å□^2^ (step 1), 5 kcal mol□^1^ Å□^2^ (step 2), 2.5 kcal mol□^1^ Å□^2^ (step 3), 1 kcal mol□^1^ Å□^2^ (step 4), 0.5 kcal mol□^1^ Å□^2^ (step 5), and 0.1 kcal mol□^1^ Å□^2^ (step 6). Each of the first three equilibration rounds lasted 500 ps, while the final three rounds lasted 1 ns each. The entire procedure of MD simulations was replicated for two additional repeats of 500 ns long simulations each, making up a total of 1.5 µs-long MD trajectory data (from triplicates).

### 2.3. MD trajectory analyses

The overall and localized structural stability of VISTA during MD simulations (from triplicates) were analyzed by assessing the evolution of VISTA’s backbone Root Mean Square Deviation (RMSD) and Root Mean Square Fluctuation (RMSF) computed using the CPPTRAJ tool from Amber Tools.^29^ The trajectories were first stripped of all TIP3P water molecules and ions, leaving only the protein and the lipid bilayer. The protein was then centered and aligned to a reference structure (initial structure) to remove any translational and rotational motions. The analysis was performed on a subset of frames from each production run. Specifically, the frames were strided by a factor of 10, resulting in one frame every 1 ns. This reduced the file size and reduced the need for high computational power. RMSD was calculated for the whole VISTA protein, and the separate segments of VISTA such as the ECD, TMD and ICD to understand the evolution and stability of the system. RMSF values were computed for the whole VISTA protein to delineate the regions of high stability and flexibility during MD simulations.

### 2.4. Principal Component Analysis (PCA)

PCA is a useful tool in MD simulations to analyze the fluctuations that occur the most in a trajectory. It accomplishes this by reducing the conformational space to a set of vectors known as principal components (PCs). Each PC captures the collective motion in the positive and negative direction of a specific vector. The first few PCs are usually able to capture the majority of the variance and fluctuations of a MD trajectory. The Bio3D package^30^ installed in R was used to perform the PCA calculations. For the non-glycosylated system, this was performed over the entire trajectory. For the glycosylated system, this was calculated between the 300-500 ns time of the trajectory. For both systems, the protein was fitted based on the C_α_ atoms to remove translational and rotational motion.

### 2.5 Modelling and simulation of the glycosylated, full-length VISTA model

The structural models for VISTA were obtained from AlphaFold.^21^ The complete structure of VISTA without the signal peptide was used to create the MD system. The cytoplasmic tail in the AlphaFold structure stretched through the transmembrane and extracellular domain so the S232-Y233 bond angle was rotated to keep it within the intracellular region. It was protonated at pH = 7.4 using the H++ server.^31^ The protein was embedded in a lipid bilayer composed of 100% 1-palmitoyl-2-oleoyl-sn-glycero-3-phosphocholine (POPC) using the CHARMM-GUI membrane builder.^32^ The initial positioning and orientation of VISTA within the membrane were determined using coordinates derived from the PPM3 (Positioning of Proteins in Membranes) method available through the PPM web server. Carbohydrates were added at N-linked glycosylation sites on VISTA (N17, N59, and N96) using CHARMM-GUI.^33^ These sites were identified based on UniProt annotations and predictions from NetNGlyc 1.0 software. The glycosylation consists of a core N-acetylglucosamine (β-GlcNac) attached to the asparagine, a second β-GlcNac linked by a β-1,4 glycosidic bond, followed by a β-mannose also attached by a β-1,4 linkage, and then two α-mannose rings that are attached to the β-mannose through α-1,6 and α-1,3 linkages. The protein system was subsequently solvated with TIP3P water molecules and ionized with NaCl.

The system was subjected to MD simulations using the GROMACS engine^34^ with the CHARMM36m force field.^35^ It was minimized and heated to 310K. It was then equilibrated in an NPT ensemble decreasing restraints in an 8-step protocol before a 500 ns production run.

## 3. Results and Discussions

### 3.1 Stability and dynamics of membrane-embedded VISTA model

Initially, a comprehensive atomistic 3D model of VISTA was constructed (as described in the Methodology section) and embedded into a solvated, electro-neutralized membrane environment (Figure 2A). This model was optimized through 500 ns long MD simulations in triplicates (totaling 1.5 μs) to gain atomic level insights into VISTA’s dynamics in its membrane-embedded state, a subject that has not been addressed previously.

**Figure 2.**
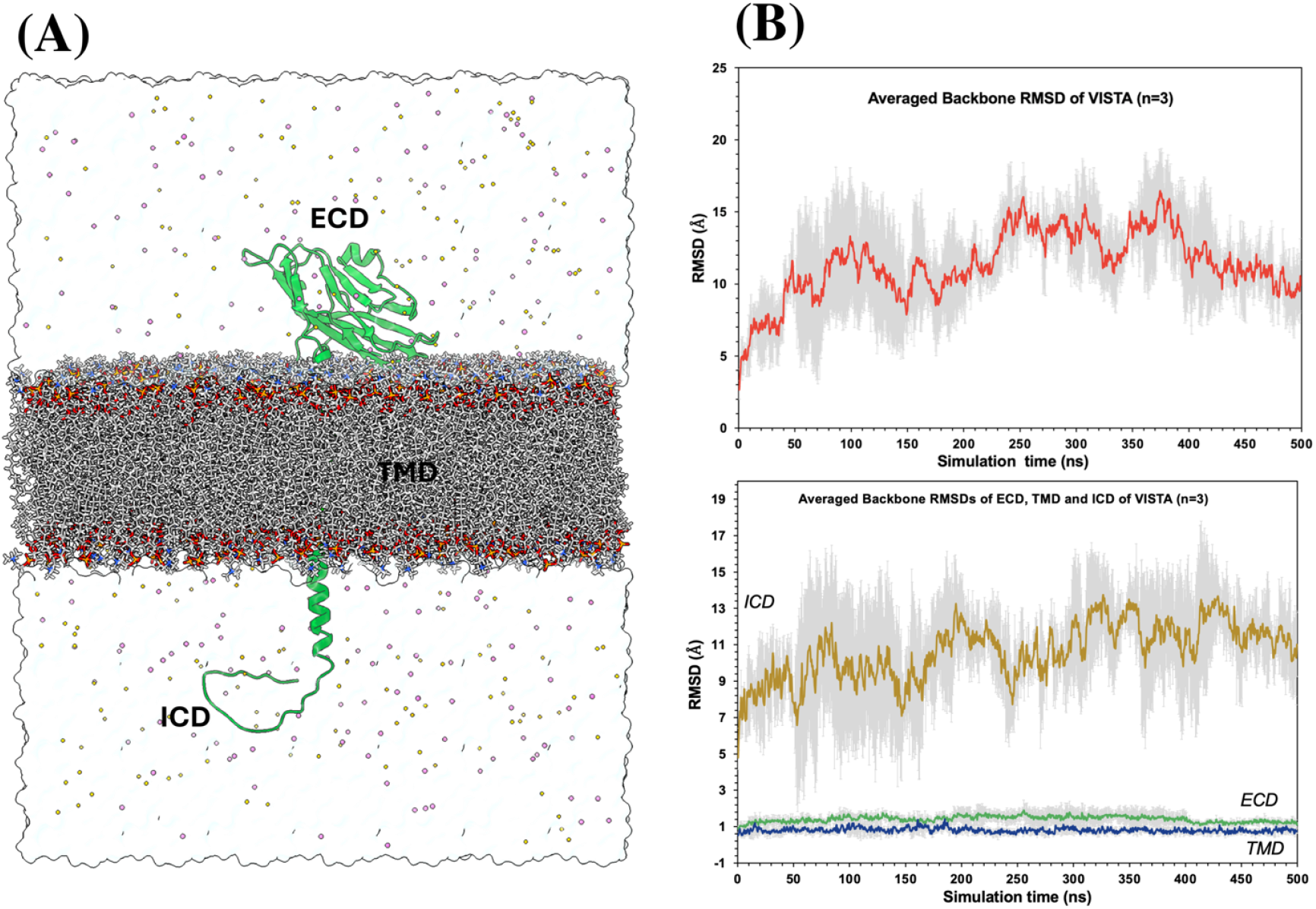
3D structure of the membrane-bound VISTA model and the evolution of backbone RMSD of the full protein and its individual domains during the MD simulations. **(A)**The atomistic model of a comprehensive VISTA structure (in green) integrated into the POPC bilayer (shown as stick representations), solvated with TIP3P water molecules (shown as a transparent white surface), and electro neutralized with counter ions (potassium ions shown as orange circles; chloride ions shown as pink circles). **(B)** The evolution of backbone RMSD is shown for the full VISTA protein (top) and its individual domains (bottom), averaged over the three MD replicates (n=3) with standard deviation. The full VISTA protein exhibited larger flexibility, predominantly attributed to the highly flexible ICD due to its inherent disordered nature. The backbone RMSD of the ECD and TMD remained highly stable during MD simulations. **Note**: In each case, the selection of residues (Proteins: residues 1-228; ECD: residues 1 to 152 (excluding the stalk segment); TMD: residues 163 to 183; ICD: residues 184 to 228) were first fitted against the initial frame and then RMSDs were calculated.

The stability of the system was evaluated by probing the evolution of the backbone RMSD of the entire protein and its independent domains such as ECD, TMD, and ICD. As shown in Figure 2B (right), the RMSD values of VISTA backbone, averaged over the MD triplicates, exhibited significant fluctuations. RMSD values increased to about 12 Å during the initial 50 ns and subsequently fluctuated between 10-15 Å. Towards the final 100 ns of simulations, the VISTA backbone RMSD plateaued at approximately 10 Å. The RMSDs of the individual domains, calculated after fitting them separately, highlighted that ECD and TMD of VISTA remained highly stable with their average fluctuations consistently below 2 Å. This confirms that the structures of these domains remained intact throughout the MD simulations. Nevertheless, ICD exhibited the largest RMSD fluctuations in the entire protein, with averaged values reaching ∼7 Å within the first 10 ns and peaking as high as 16 Å. This RMSD behavior correlated with the overall RMSD changes for the full VISTA protein. Similar RMSD trends were observed in each of the individual MD repeats (Supplementary Information, Figure S3), confirming that the ECD and TMD of the VISTA reached convergence while ICD contributed majorly to the overall variability.

The stability of VISTA’s internal structure was further assessed by calculating the evolution of its Radius of Gyration (*R*_*g*_), a measure for the compactness of the system during MD simulations. The *R*_*g*_ values of ECD and TMD, the core segments of VISTA, remained relatively stable confirming their compactness and well-folded structures (Supplementary Information, Figure S4). Nevertheless, the Rg values of the ICD showed large variations, highlighting its high instability during simulations.

To further confirm this aspect and delineate the stable and flexible parts of VISTA, per-residue backbone RMSF was analyzed after aligning the complete VISTA model. The averaged RMSF plot of VISTA (in Figure 3A) agree well with the inference from the RMSD and Rg analyses. The most stable RMSF values were observed for the TMD residues, while the largest fluctuations were observed for the residues making up the ICD. This is unsurprising due to the highly disordered nature of this domain in VISTA. Bioinformatics programs employing deep learning and machine learning, such as AIUPred^36^ and Pondr^37^ (with XL-VRT variant), were used to predict the disordered regions in the VISTA sequence. Both programs consistently predicted the residues that form the VISTA ICD to be highly disordered (a prediction score of > 0.5 on a scale of 0-1), as shown in Figure 3B. Previous studies^1^ have reported that the VISTA ICD interacts with different types of proteins. For example, a four-residue motif, N^210^-P^211^-G^212^-F^213^, in VISTA’s ICD have been shown to interact with the adapter protein, NUMB, which promotes its checkpoint signaling.^38^ VISTA ICD has also been confirmed to interact with galectin-9 inside the cell.^39^ In addition, the cytoplasmic tail of VISTA ICD also features a number of potential phosphorylation sites for kinases such as MAPKs.^40^ Thus, the disorderly nature of VISTA is plausibly suited to its variable roles and may become more ordered upon protein-protein interactions inside the cells.

**Figure 3.**
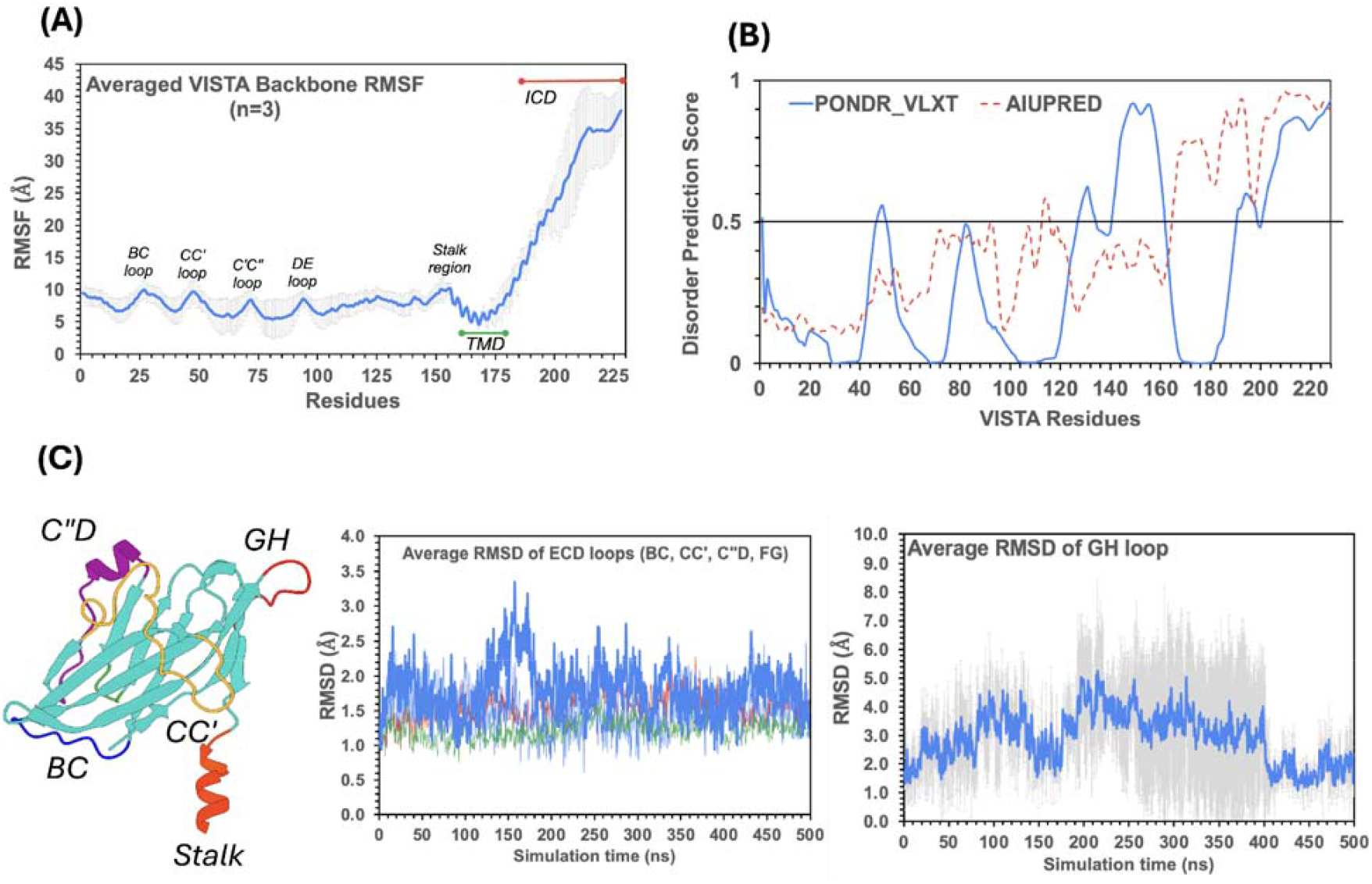
Per-residue RMSF analyses of VISTA, its ICD disorder prediction along with ECD loop dynamics. (A) The averaged backbone RMSF of VISTA highlights that the ICD was the most dynamic segment of the protein. (B) AI-based predictions (using PONDR and AIUPreD programs) based on the amino acid sequence of VISTA (residues 1-228 employed in this work) confirm the disordered nature of the ICD, explaining its flexibility during MD simulations. (C) Most of the loops in the VISTA ECD remained stable, except for the GH loop, which exhibited larger flexibility. The GH loop connects the last beta strand that is unique to VISTA.

Nevertheless, the VISTA ECD remained highly stable. It was interesting to note that most of the ECD loops, which are usually expected to be flexible, remained significantly stable (Figure 3C). All the loops connecting the BC strands, CC’ strand, C”D strand, and the FG strands, critical in promoting VISTA’s interactions with its ligands (VSIG-3 and PSGL-1) and monoclonal antibodies, remained stable with a fluctuation less than 3 Å in all the MD repeats. Especially, the longest CC’ loop in the VISTA ECD was particularly highly rigid during MD simulation. This is consistent with the behavior of apo VISTA ECD in our earlier work.^1^ This stability can be attributed to the C51^CC’loop^-C113^F-strand^ disulfide bond that connected this loop with the F β-strand of VISTA ECD (see in Figure 1C). In addition, there were a number of hydrogen bonds (H-bonds) that supported the stability of CC’ loop, as described in the next section. Nevertheless, the GH loop, despite involving the C146^GH loop^-C12^A-strand^, displayed a very large flexibility during the MD simulations (Figure 3C). Such higher fluctuations of GH loop have also been reported by a previous MD-based studies of VISTA ECD.^1, 40^ The H-strand and its preceding GH loop are involved in VISTA’s interactions with its ligands and hence remains flexible in the absence of a ligand partner. Furthermore, the stalk region of VISTA (residues 153-162) connecting the ECD with that of the TMD exhibited higher fluctuations, which supported the dynamics of ECD.

### 3.2 Interactions promoting unusual stability of CC’ loop of VISTA

The CC’ loop of IgV domains in immune checkpoint proteins is a key determinant of ligand recognition at the immunological synapse and contributes directly to T-cell inhibitory signaling. In most checkpoints, this loop is highly flexible in the apo state and undergoes conformational rearrangement upon ligand engagement.^41-46^ For example, the CC’ loop of apo PD-1 exhibited substantial deformation during MD simulations and adopted a distinct conformation when interacting with PD-L1.^41-44^ Similar flexibility has been reported for CTLA-4^46^ and CD48.^45^ Notably, in these checkpoints, the CC′ loop is relatively short and typically made up of 5-7 residues (four residues in CTLA-4 and seven in PD-1). In contrast, the CC’ loop of VISTA (residues S43–H66), which is unusually long at ∼24 amino acids, displayed exceptional rigidity in our apo simulations, with average RMSD fluctuations below 2.5 Å (Supplementary Information, Figure S5). While the C51^CC’loop^-C113^F-strand^ disulfide bond covalently connected the CC’ loop with the F-strand, our analyses revealed a dense network of H-bonds between the CC’ loop and the surrounding regions that collectively contributed to the overall stability of this loop in VISTA.

Figure 4 presents both qualitative and quantitative analyses of H-bonds involving residues of VISTA’s CC′ loop. Approximately 11 of the 24 residues (∼46%) in the CC′ loop participated in H-bond interactions, with most interacting pairs exhibiting distances of ∼2-3.5 Å (Figure 4A). In our previous study of the apo VISTA ECD^1^, an arginine “fingerprint” region was identified as forming multiple H-bonds; these interactions were also observed in the full-length VISTA model, highlighting their critical role in stabilizing the CC′ loop.

**Figure 4.**
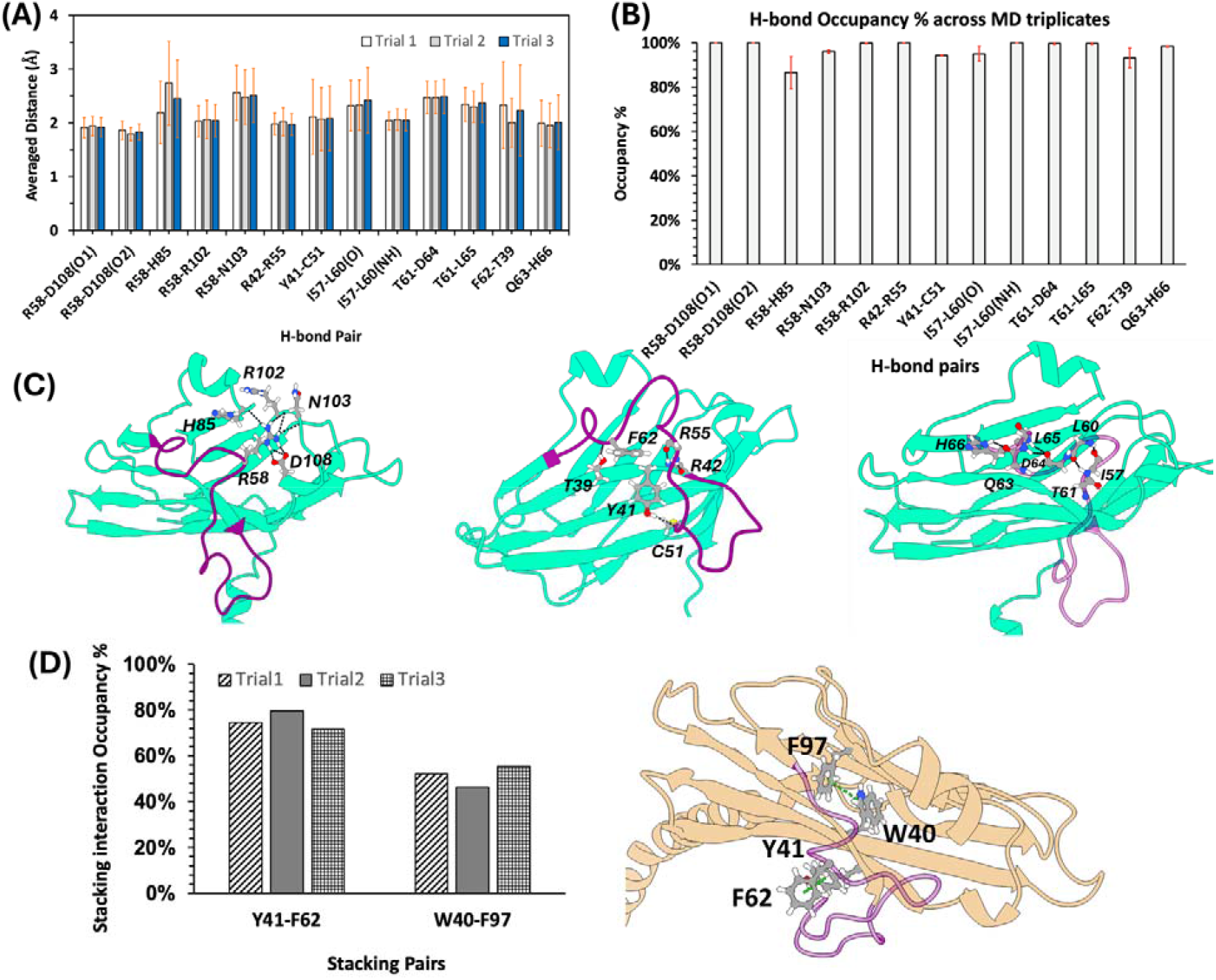
Quantitative analysis and structural mapping of key molecular interactions in VISTA ECD. (A) The averaged heavy-atom H-bond distances for the defined interaction pairs plotted across the three independent MD simulations. (B) The corresponding H-bond occupancy percentages, averaged over the replicates, further support the stability of these interactions. (C) Representative structural views map these contacts onto the protein, with the ribbon shown in cyan, interacting residues depicted in atomic detail, the CC′ loop highlighted in purple, interaction pairs shown as ball-and-stick models, and the hydrogen bonds illustrated as black dashed lines. Time evolution data of individual H-bonds are provided in supplementary information, Figure S6. (C) Aromatic stacking interactions between Y41^C-strand^-F62^CC’loop^, and W40^C-strand^-F97^D-strand^ residue pairs in VISTA ECD (Left). Both pairs involved in an intermediate T-shaped stacking interaction (Right). 3D structure of VISTA ECD (in yellow cartoon), and the aromatic stacking pairs shown as ball and stick representations. The Y41^C-strand^-F62^CC’loop^ was exposed to the surface, while the W40^C-strand^-F97^D-strand^ was present within the central hydrophobic core of VISTA.

As illustrated in Figure 4A-C, the sidechain of R58^CC’ loop^ was oriented towards a central hydrophobic core of VISTA and engaged in multiple H-bonds with residues such as H85^〈2 helix^ (backbone), and R102 (backbone), N103 (backbone), and D108 (sidechain) from EF loop (Figure 4A-B). The occupancy percentages, defined as the fraction of frames with a donor-acceptor distance < 3.5 Å, averaged over the MD triplicates, were 100% for most of these interactions, while the R58-H85 pair showed a slightly reduced occupancy of ∼85%. In addition, the backbone atoms of R55, C51, and F62 within the CC′ loop formed stable H-bonds with R42 (backbone), Y41 (sidechain), and T39 (sidechain), respectively, from the C strand of VISTA. These interactions also consistently remained below the 3.5 Å threshold, with average occupancies exceeding 90%. Apart from its interactions with surrounding structural elements, the CC′ loop also formed several stabilizing intra-loop H-bonds. Backbone-backbone H-bonds were observed between residue pairs such as I57-L60, T61-D64, and T61-L65, involving their respective amino and carboxyl groups. Additionally, the backbone carboxyl group of Q63 formed a H-bond with the sidechain imidazole of H66. These intra-CC′ loop interactions were also highly stable, consistently remaining within the 3.5 Å distance cutoff and exhibiting occupancies greater than 90%. These observations underscore the importance of this H-bond network in stabilizing the CC′ loop and, more broadly, maintaining the structural integrity of the VISTA ECD. The time evolution of H-bond distances for the individual pairs across all the MD replicates are provided in Supplementary Information, Figure S6.

Beyond H-bonding, the structural integrity of the CC’ loop is further reinforced by a persistent aromatic stacking interaction between F62^CC’loop^ and Y41^C-strand^. This interaction adopts an intermediate T-shaped geometry and remains highly stable throughout the simulations, maintaining an occupancy of 72–80% across all MD replicates (Figure 4D and Supplementary Information, Figure S7). The high stability of this stacking pair is functionally significant, as it preserves the optimal surface exposure of F62. This residue has been identified as a critical component of the three-amino acid epitope (R54/F62/Q63) responsible for VSIG-3 binding^47^, and is also known to facilitate essential aromatic interactions with mAbs.^1, 13, 14^ By anchoring the CC’ loop in a favorable conformation, this stacking interaction ensures that the VISTA interface remains competent for both therapeutic targeting and biological ligand recognition. In contrast to the surface-exposed interaction, a second aromatic stacking pair was identified between W40^C-strand^ and F97^D-strand^, oriented towards the central hydrophobic core of VISTA (Figure 4D). This internal interaction was found to be relatively weaker and more dynamic, with occupancy levels ranging between 46% and 55% across the trials. The lower persistence of this buried pair suggests a degree of internal plasticity within the hydrophobic core, whereas the surface-exposed F62-Y41 interaction provides the necessary rigidity to maintain the protein’s functional binding interface.

### 3.3. Conformational dynamics of VISTA

PCA and conformational clustering were performed independently on each of the three MD replicates to characterize the conformational dynamics of VISTA during simulations (Figure 5). In each case, the high-dimensional MD trajectory data were projected onto a reduced-dimensional conformational space, allowing dominant collective motions to be identified while minimizing contributions from high-frequency, uncorrelated fluctuations. This analysis highlights the essential dynamics of VISTA, i.e., the motions that are most likely to be relevant to its structure and function. The PCA scree plots (Figure 5, right column) revealed that VISTA’s conformational variance is dominated by a limited number of collective modes. In all three replicates, the first two principal components (PC1 and PC2) together accounted for the majority of the total variance, capturing between ∼52% and ∼75% of the overall motion. Notably, in replicate 3, PC1 alone accounted for ∼64% of the variance, indicating that VISTA’s dynamics are largely governed by a small number of dominant, low-dimensional collective motions rather than being evenly distributed across many degrees of freedom.

**Figure 5.**
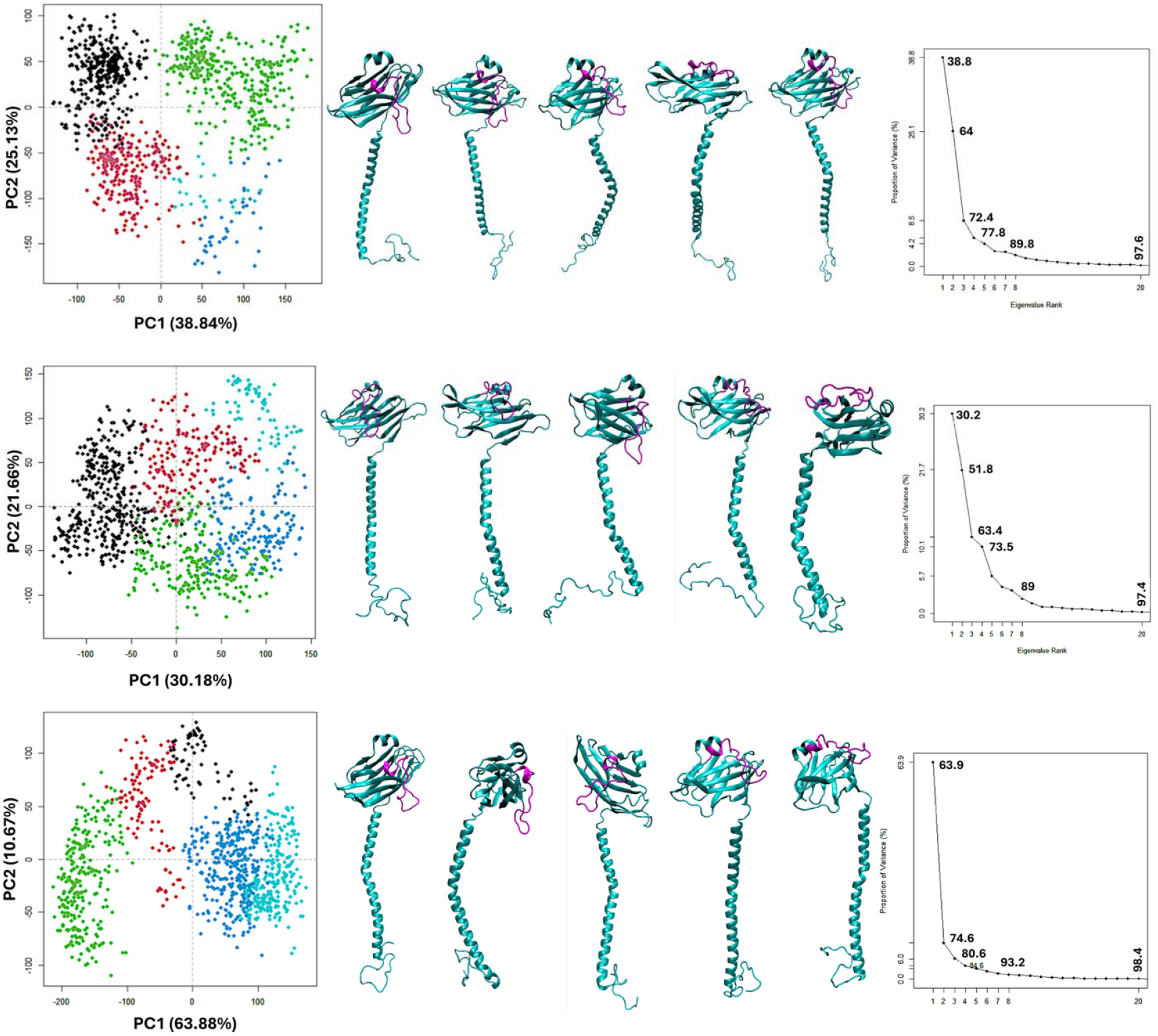
PCA and conformational clustering of VISTA essential dynamics across independent MD replicates. The conformational landscape of VISTA is explored through the 2D projection of trajectories onto the first two principal components (PC1 and PC2) for three independent simulations (Repeat 1, top; Repeat 2, middle; and Repeat 3, bottom). Distinct clusters in PCA plots are color-coded to represent the sampling of metastable states. Representative structures from each cluster are shown (Middle Column) with the CC’ loop colored in purple and the remainder of the protein depicted in cyan ribbon. These snapshots reveal two distinct, coupled motions: (i) a coordinated rotation of the ECD that modulates the positioning of the CC′ loop, transitioning it between a membrane-dipping orientation facing the TMD interface and an upward-facing conformation; and (ii) the bending of the TMD to accommodate these ECD changes. The magnitude of these essential motions is quantified via scree plots (Right Column), which illustrate the proportion of variance captured per eigenvalue rank. The dominance of the first few components underscores that the large-scale bending and twisting of the ECD-TMD linker constitutes the primary functional motion of the system.

The 2D projection of PC1 versus PC2 from the MD trajectories revealed that VISTA explored a broad conformational space, indicative of a rugged free-energy landscape. To identify metastable states sampled during the simulations, the conformational ensembles were clustered into five distinct groups. Clear spatial separation between the clusters, particularly in repeats 1 and 3, was observed, suggesting the presence of significant conformational transitions between well-defined structural states. To further characterize these metastable states, representative snapshots from each cluster across the three MD replicates were extracted (Figure 5, middle column). Visual inspection of these representative conformations revealed that the VISTA ECD underwent substantial rotational motion relative to the membrane normal. This rotation led to pronounced modulation of the CC′ loop orientation. In a certain conformational state (dubbed as a closed or down conformation in this text), the CC’ loop was positioned downward toward the TM domain, enabling it to interact with the lipid bilayer. Whereas, in other conformational states (Open or up conformation), the ECD was rotated such that the CC’ loop was projected away from the TM domain and remained largely disengaged from membrane contacts.

To quantitatively analyze the membrane interactions of the CC’ loop in the down (or closed) conformation, contact analyses of the CC’ loop residues 43-56 with the POPC bilayer were carried out (Figure 6). For this purpose, the distance between each of the residues (any atoms) and any of the POPC atoms was computed with a measuring distance capped at 15 Å. A threshold of 5 Å distance was applied to consider any kind of non-bonded interactions between the loop and membrane, and their occupancy % was estimated. Across all three MD replicates, residues 47-50 consistently exhibited notable contact occupancy of over 20% (Figure 6, top panel), indicating that this central region of the CC′ loop functions as a key membrane-anchoring hotspot. Other flanking residues (43-46 and 50-51) also occasionally contacted the bilayer, but with lower occupancy, suggesting a dynamic anchoring mechanism where the central residues stabilize membrane engagement while the surrounding residues remain flexible.

**Figure 6.**
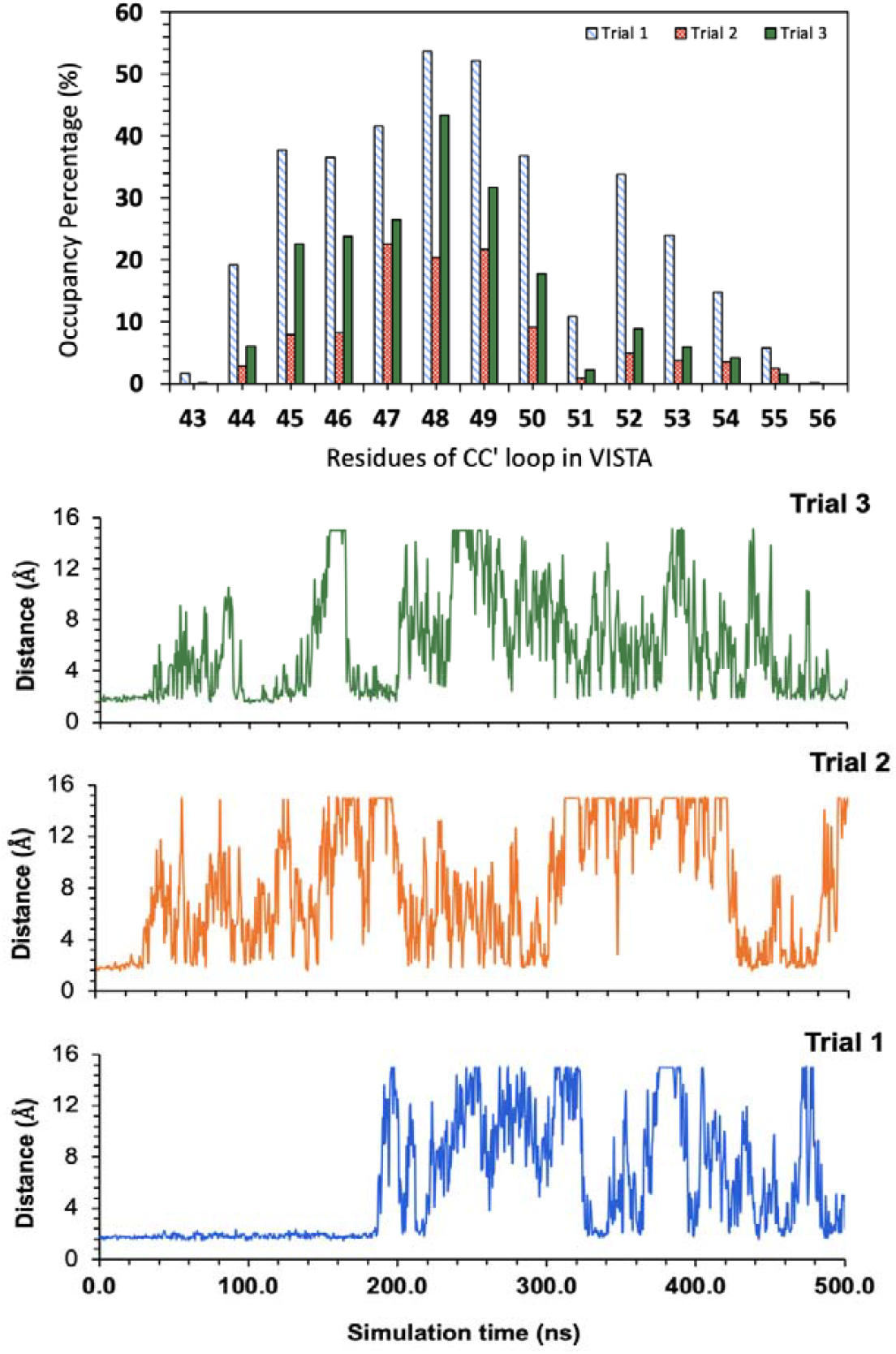
Membrane interactions of the VISTA CC′ loop across independent MD replicates. The interaction between the CC′ loop of VISTA (residues 43–56) and a POPC bilayer was quantified across three independent MD simulations. **Top:** Residue-wise membrane contact occupancy, expressed as the percentage of simulation time each residue remains within 5 Å of any POPC atom. Residues in the central region (47-49) consistently exhibit high occupancy, identifying them as key membrane-anchoring hotspots. **Bottom:** Time evolution of the minimum distance between POPC and the CC′ loop residues with high occupancy (43-50). For each frame, the shortest distance between any atom of these residues and any POPC atom was calculated. Distances were capped at 15 Å for visualization. The trajectories reveal frequent membrane engagement events (<4 Å), highlighting dynamic membrane interactions with specific residues acting as transient yet recurring anchors to the lipid surface.

Time-resolved analysis of the minimum distances between the CC′ loop and the membrane (Figure 6, bottom panels) revealed frequent events where the loop approached the bilayer to < 5 Å, confirming that these interactions are transient yet recurrent. Notably, all three trials showed a similar pattern of dynamic engagement, with the loop repeatedly approaching and retracting from the membrane over hundreds of nanoseconds, highlighting the flexible yet functionally relevant nature of the CC′ loop in mediating VISTA-membrane contacts. Such dynamic membrane contacts of the CC’ loop highlight the rotation of CC’ loop, sampling between the open (CC’ loop away from the membrane) and closed (CC’ loop interacts with POPC) states throughout the MD simulations, as identified by the PCA and clustering analyses (in Figure 5).

The ECD rotation in VISTA is allosterically supported by the bending of its long TM helix, which is facilitated by a proline residue (P177) in TM domain. Proline-induced bending of TM helices has been widely reported in integral membrane proteins^48-52^ and often plays critical roles in gating, conformational switching, and structural adaptability. Notably, while a few other immune checkpoints, such as CD80, CD86, PD-L2, and ICOS (in Supplementary Information, Figure S8), contain TM prolines, the TM helical bending and ECD rotation to mediate membrane interactions has not been reported in any of these proteins. X-ray crystallography and biochemical binding assay of murine TIM-4 protein showed that the protein interacts with phosphatidylserine (PDB 3BIB), particularly through its CC’ loop, along with other loops.^53^ Indeed, the authors have provided a schematic image, hypothesizing that the ECD of mTIM would bend downwards to allow its CC’ loop interaction with the phosphatidylserine.^53^ This proposed mechanism is similar to the membrane-dipping behavior observed for VISTA ECD in our MD simulations.

### 3.4. ECD rotation observed in glycosylated VISTA model

Although the MD simulations of our VISTA model were performed in triplicate to ensure reproducibility, simulations can still be influenced by the starting configuration and the specific MD program used. Furthermore, our initial model lacked glycosylation, whereas immune checkpoint proteins are typically glycosylated, which could impose steric or dynamic restraints on the ECD and modulate its rotation. To test the ECD rotation hypothesis under more physiological conditions, we constructed a comprehensive, glycosylated, membrane-integrated model of human VISTA based on the AlphaFold^21^ structure. The ICD in the AlphaFold model was misoriented and would have artificially inserted into the membrane; to correct this, the ICD was rotated by altering the S232-Y233 bond angle to prevent membrane interactions. Subsequently, three asparagine residues (N17, N59, and N96) were modified with high-mannose type N-glycans, and the fully glycosylated model was embedded into a POPC bilayer using the CHARMM-GUI server. This model (Figure 7A) was subjected to a 500 ns MD simulation using GROMACS^34^ and the CHARMM36m force field^35^ to evaluate whether glycosylation constrains ECD rotation.

**Figure 7.**
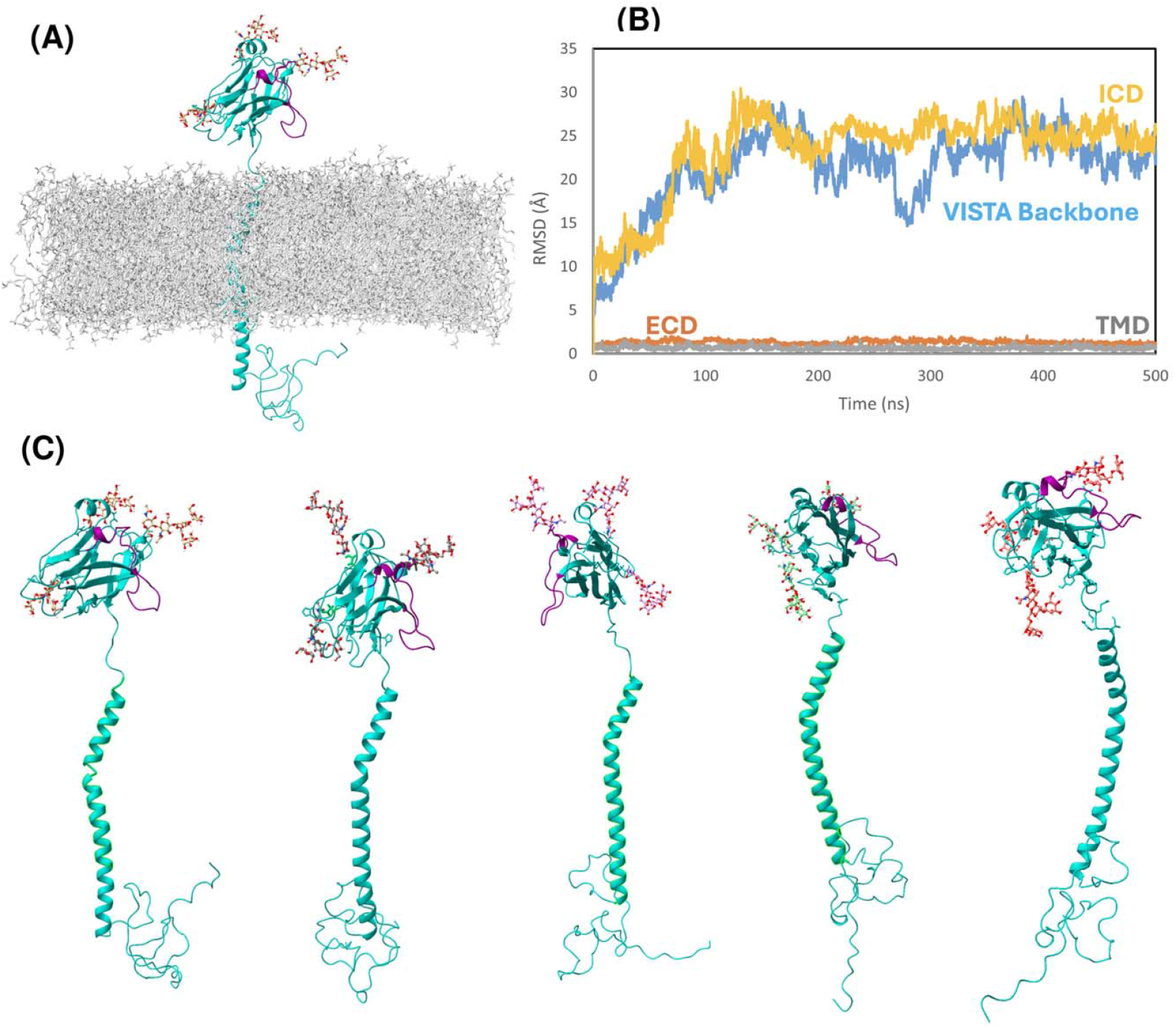
Conformational dynamics of the glycosylated, membrane-bound, full-length VISTA model. (A) The 3D structure of full-length VISTA, with the protein shown in cyan cartoon representation, the CC′ loop in purple, glycans in ball-and-stick representation, and the lipid bilayer in grey sticks. (B) Time evolution of backbone RMSD shows that VISTA initially fluctuated, reaching a maximum deviation of ∼25 Å during the first 200 ns, after which fluctuations stabilized within ∼5 Å. The ICD exhibited the largest fluctuations, confirming its dynamic nature, whereas the ECD and TMD remained highly stable, with RMSD <3 Å throughout the simulation. (C) Representative snapshots from five clusters identified in PCA analyses (Supporting Information, Figure S9) illustrate the rotation of the ECD. This rotation positions the CC′ loop either toward the TMD, allowing interactions with the membrane, or away from the TMD, remaining largely disengaged from the membrane. Overall, the RMSD profiles and structural dynamics of the full-length, glycosylated model closely resemble those observed in the truncated, glycan-free VISTA model.

Initially, the backbone RMSD of the full-length, glycosylated VISTA model was calculated over the 500 ns MD simulation (Figure 7B). The overall RMSD profile closely resembled that observed for the glycan-free, truncated VISTA model (Figure 2), indicating that the inclusion of glycans did not substantially alter the global dynamics of the protein. During the first ∼200 ns, the backbone RMSD increased gradually, reaching a maximum deviation of ∼25 Å, primarily due to large-scale motions of the ICD. After this initial equilibration phase, the RMSD stabilized within ∼5 Å for the remainder of the simulation, demonstrating convergence of the overall structure. When analyzed by domains, the ECD and TMD exhibited remarkable stability, maintaining RMSD fluctuations below ∼3 Å throughout the simulation, consistent with their compact and well-folded conformations. In contrast, the ICD displayed larger deviations during the early phase of the simulation, reflecting its intrinsically disordered and flexible nature.

Further, PCA analyses were performed on the MD trajectory of the full-length model to delineate its essential dynamics and confirm if the up (open) and down (closed) conformational states were sampled. Initially, the scree plot from PCA (Supporting Information, Figure S9) revealed that the first five PCs collectively accounted for ∼89% of the total variance, with PCs 1 and 2 alone capturing over 50% of the variance. The dominance of these low-dimensional modes once again indicates that the protein’s motion is not random but largely governed by coordinated, large-scale conformational transitions. Projection of the trajectories onto the PC1 Vs PC2 space identified distinct clusters (color-coded in red and blue), representing metastable states sampled during the simulations.

Representative snapshots from these clusters (Figure 7C and Supporting Information, Figure S9) illustrate that the primary functional motion is a substantial rotation of the ECD relative to the membrane normal. This rotation modulates the orientation of the CC’ loop (purple), giving rise to two dominant conformational states. The Down state, where the CC’ loop rotated toward the TMD, facilitating interactions with the lipid bilayer; whereas, in the Up conformation, the ECD rotated away from the membrane, leaving the CC’ loop disengaged and accessible for extracellular ligand binding. It is interesting to note that the N-glycans attached at N17, N59, and N96, did not sterically hinder this rotation. Thus, the conformational landscape sampled by the glycosylated model closely mirrored that of the truncated, glycan-free system. This agreement confirms that ECD rotation is an intrinsic mechanical property of VISTA, likely critical for its function as a dynamic immune checkpoint regulator. Glycosylation may act more subtly, potentially modulating the kinetics and frequency of transitions between the up and down states rather than acting as a rigid structural anchor.

### 3.5 Potential functional implications of VISTA Up/Down conformations

It is well-established that the CC’ loop of IgV domains from the immune checkpoint proteins play a critical role in mediating the cell surface protein-protein recognition at the immunological synapse.^45^ This is also confirmed in the case of VISTA, where the CC’ loop promotes the interaction with its protein partners such as VSIG-3 and PSGL-1. Indeed, antibodies designed to target the CC’ loop of VISTA has been demonstrated to reverse immune suppression and inhibit tumor growth.^10^ Therefore, accessibility to CC’ loop is critical for VISTA’s cognate partners to interact with it and suppress immune suppression.

Our MD simulations reveal that VISTA’s extracellular domain (ECD) undergoes significant rotational motions relative to the membrane, sampling transient “Up” and “Down” conformations that reorient the CC’ loop. While other ECD loops (e.g., FG and BC) also contact the membrane during these dynamics, the CC’ loop exhibits the most pronounced conformational changes. In the Up conformation, the CC’ loop is fully accessible for potential ligand binding. In the Down conformation, the loop engages in membrane interactions. Our non-bonded interaction analyses (in Figure 4) revealed that ∼46% of residues making up the CC’ loop were involved in highly stable H-bond interactions, in addition to a couple of stacking interactions. Only a short stretch of CC’ loop residues (43-50) were not involved in persistent molecular contacts. Interestingly, this short stretch is the main segment that interacted with the membrane bilayer when VISTA sampled the “Down” conformation. Therefore, we hypothesize that the VISTA may undergo this Up/Down conformational switching mechanism to regulate its immune checkpoint signaling.

Classically, immune checkpoint receptors engage ligands on neighboring cells (*trans* interactions), for example, PD-1 on T cells binding to PD-L1 on APCs. Until recently, this model was assumed for VISTA interactions with VSIG-3 and PSGL-1. However, cell-cell aggregation experiments indicate that membrane-bound VISTA does not mediate detectable trans interactions with VSIG-3 or PSGL1^54, 55^ under physiological or acidic pH conditions. Instead, complementary assays demonstrate that VISTA engages these ligands in a *cis* configuration, on the same cell surface.^59^ To facilitate such cis interactions, VISTA may require substantial ECD reorientation, consistent with the rotational dynamics observed in our simulations. The bending of the TM helix, enabled by the presence of a proline residue, likely supports these conformational transitions, providing an intrinsic mechanism that enables VISTA to engage ligands selectively and modulate immune suppression. Notably, the Up conformation exposes the CC’ loop, suggesting that trans interactions, while not yet observed, cannot be fully excluded under specific cellular contexts.

## 4. Conclusion

This work focused on understanding the structure and dynamics of human VISTA in a membrane environment. A glycan-free truncated model of hVISTA (ICD truncated after R228) was embedded in a POPC bilayer and subjected to 1.5 µs of MD simulations (500 ns per replicate). Our analyses revealed that the ECD and TMD remained highly stable, while the ICD was highly flexible due to its disordered nature. Notably, the unusually long (∼24 amino acids) CC′ loop remained largely stable, supported by extensive H-bonds, aromatic stacking, and a disulfide bond involving ∼46% of its residues.

PCA and cluster analyses showed that, although the ECD is generally stable, it undergoes rotational motions sampling two distinct conformations: Up and Down. These motions reposition the CC′ loop either away from the membrane (Up state), making it accessible for ligand binding, or toward the membrane (Down state), promoting transient membrane interactions. These dynamic conformations were also reproduced in a glycosylated, full-length hVISTA model, suggesting that ECD rotation is an intrinsic property of the protein.

The Up/Down switching likely underpins VISTA’s functional regulation, facilitating cis interactions with ligands such as VSIG3 and PSGL-1 on the same cell surface. The Up conformation, exposing the CC′ loop away from the membrane, may also allow trans interactions in specific cellular contexts, although this remains to be experimentally demonstrated. Overall, these MD simulations provide novel mechanistic insights into VISTA dynamics and open avenues for the rational design of checkpoint-targeted cancer immunotherapeutics.

## 5. Author contributions

AG acquired funding to support this work, conceived the idea, supervised the work, wrote the manuscript. VR performed all the modelling, simulation, and analyses of the glycan-free model of VISTA; NL performed the modelling, simulation and analyses related to the glycosylated, full-length VISTA. AG contributed to analyzing the results, and generating the final figures reported in the manuscript, with help from VR and NL.

## 6. Declaration of Competing Interest

The authors declare that they have no known competing financial interests or personal relationships that could have influenced the work reported in this manuscript.

## 7. Acknowledgements

This research is supported by an operating fund provided to AG by Grant 946058 from the Cancer Research Society Charlotte Légaré Memorial Fund. All computational modelling and simulation presented in this work were carried out using the High-Performance Computing resources from Digital Research Alliance of Canada (https://alliancecan.ca/) made available through a Resource Allocation Competition.

## 8. Supporting Information

Supporting Information: AlphaFold model, the truncated VISTA model, time evolution data for backbone RMSD of different segments of VISTA, radius of gyration, H-bonds, and aromatic stacking interaction parameters across different MD repeats, PCA data and representative snapshots of glycosylated, full-length VISTA model.

## 9. Data and Software Availability

The computational models were built using AlphaFold [https://alphafold.ebi.ac.uk/] and CHARMM-GUI [https://www.charmm-gui.org/]. Molecular dynamics simulations and trajectory analyses were performed using widely used programs: NAMD 3.0.1, GROMACS 2022.3, VMD 1.9.3, UCSF Chimera 1.9. All procedures employed in this work are described in the methods section. The data files related to the plots in this work are provided as the Supporting Information.

## Notes

### Competing Interest Statement

The authors have declared no competing interest.

