## Supplementary Information, Figure S1 for "Membrane-coupled conformational switching of VISTA regulates immune checkpoint signaling via CC′ loop accessibility"

*Corresponding author:

Aravindhan Ganesan,

^#^ Equal first authors

**Figure S1.**

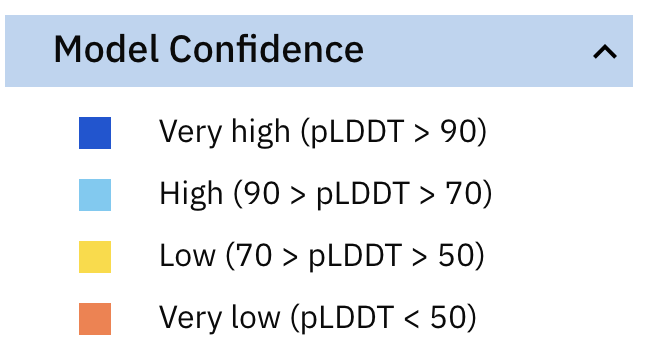

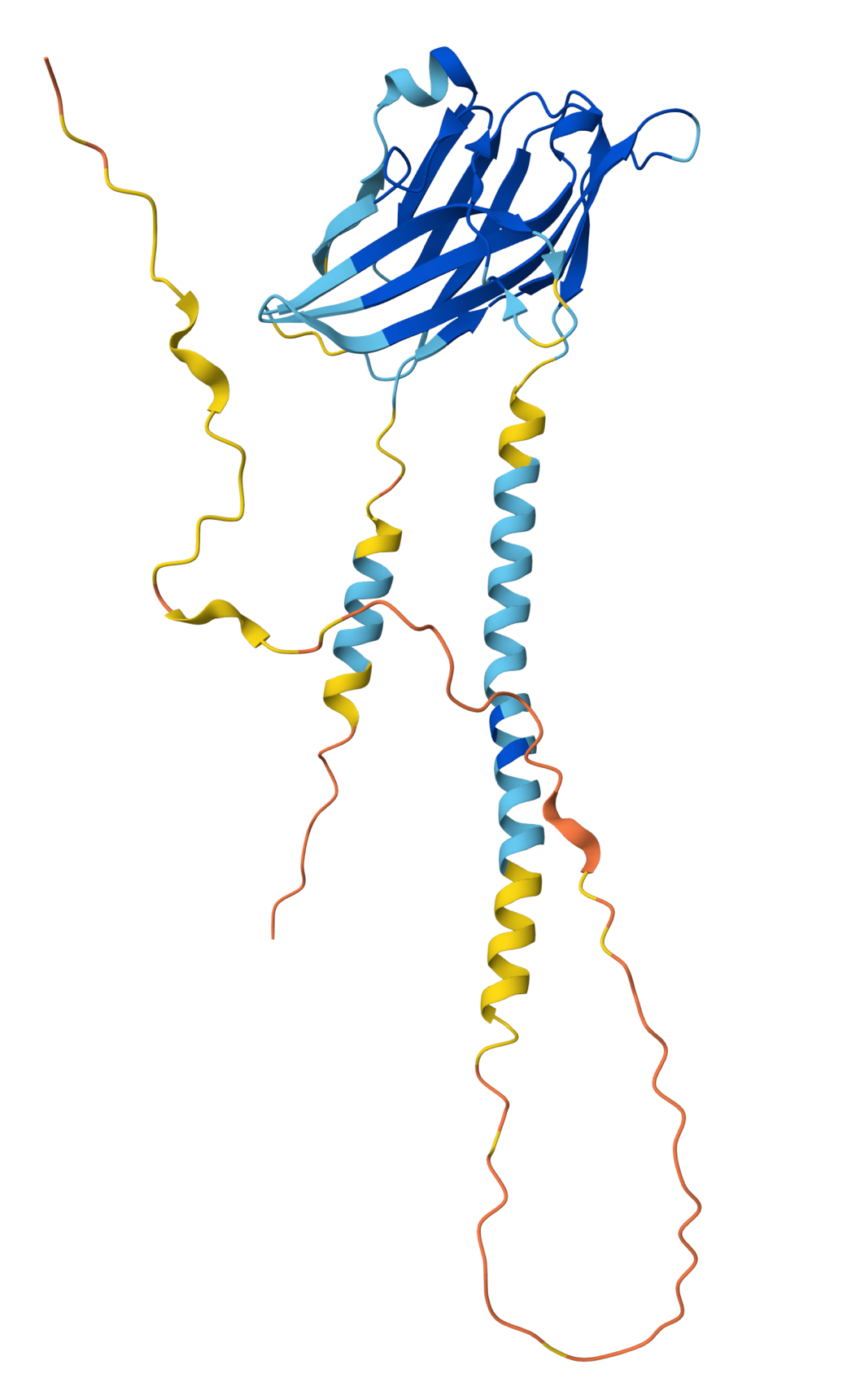

**Figure S1: The 3D structure of the full-length human VISTA modelled by AlphaFold (AF-Q9H7M9-F1-v4)** **along with the predicted model confidence.** AlphaFold has predicted the complete structure of VISTA (shown as cartoon representations) including its N-terminal signal peptide (of 32 residues), TMD, and the ICD segments. Given the availability of the experimental ECD structures, unsurprisingly, AlphaFold has predicted the ECD structure with very high to high confidence, as confirmed by the predicted local distance difference test (pLDDT) score in the range 70-90. While the TMD has also been predicted with moderate confidence (pLDDT > 50), the majority of the VISTA ICD was modelled with very low confidence (pLDDT < 50). Further, the ICD of VISTA was completely disordered in nature and is also misoriented towards TMD and ECD, making it difficult to integrate this model in the membrane bilayer.

**Figure S2**

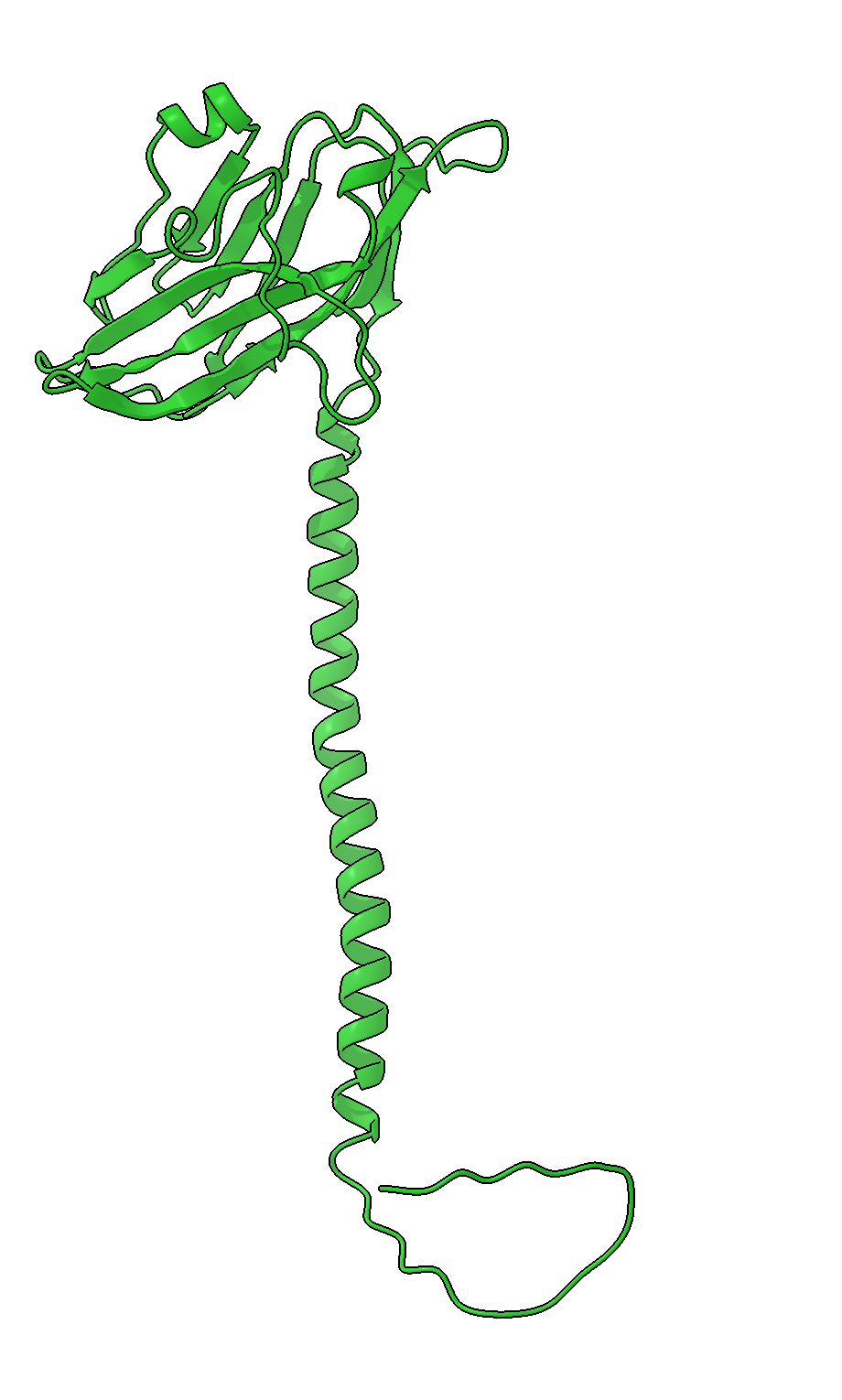

**Figure S2: The final VISTA model employed in this work.** This model (shown in green cartoon) is modelled by replacing the ICD from the AlphaFold structure (AF-Q9H7M9-F1-v4) with a truncated ICD structure remodeled through ab initio approach. The truncated ICD, comprising residues Y184-R228, was generated using Rosetta *ab initio* modeling. This comprehensive model was used as the starting point for all MD simulations and analyses in this work.

**Figure S3**

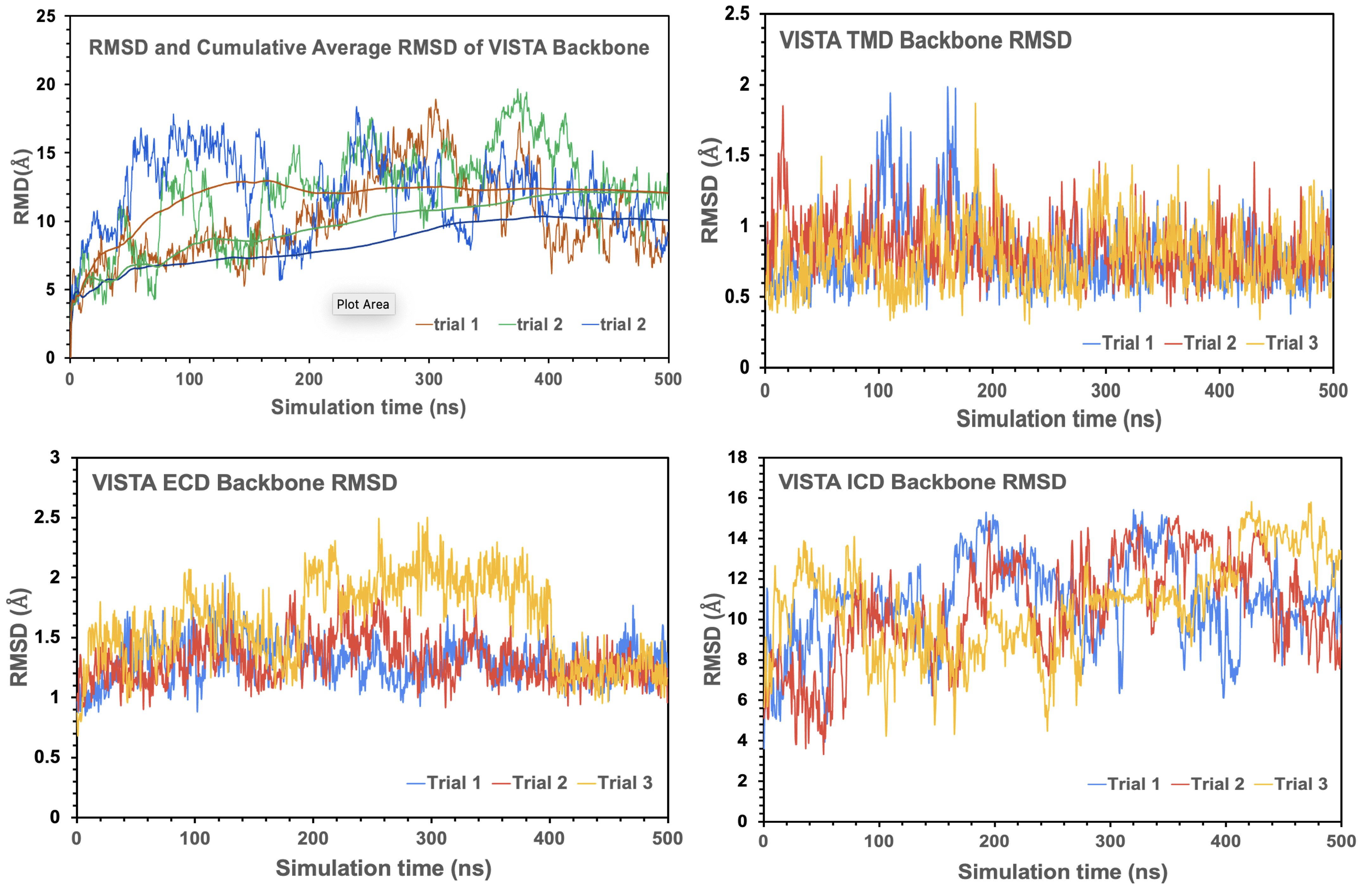

**Figure S3. The evolution of Backbone RMSD of VISTA and its individual domains during the MD simulations (triplicates).** In each case the selection of residues (Proteins: residues 1-228; ECD: residues 1 to 152 (excluding the stalk segment); TMD: residues 163 to 183; ICD: residues 184 to 228) were first fitted against the initial frame and then RMSDs were calculated. **Left Panel:** Evolution of protein backbone RMSD during the triplicate 500-ns MD simulations, along with the cumulative Average of RMSDs (Top) describes the flexibility and internal dynamics of VISTA. However, the backbone RMSD of the ECD (Bottom) exhibited larger stability with fluctuations < 2 Å during most parts of the triplicate MD simulations. **Right Panel:** The evolution of the backbone RMSD of the TMD (Top) exhibited highly stable behavior, while the backbone RMSD of the VISTA ICD exhibited the largest fluctuations in the protein during MD simulations, highlighting its flexibility due to its disorderly nature.

**Figure S4**

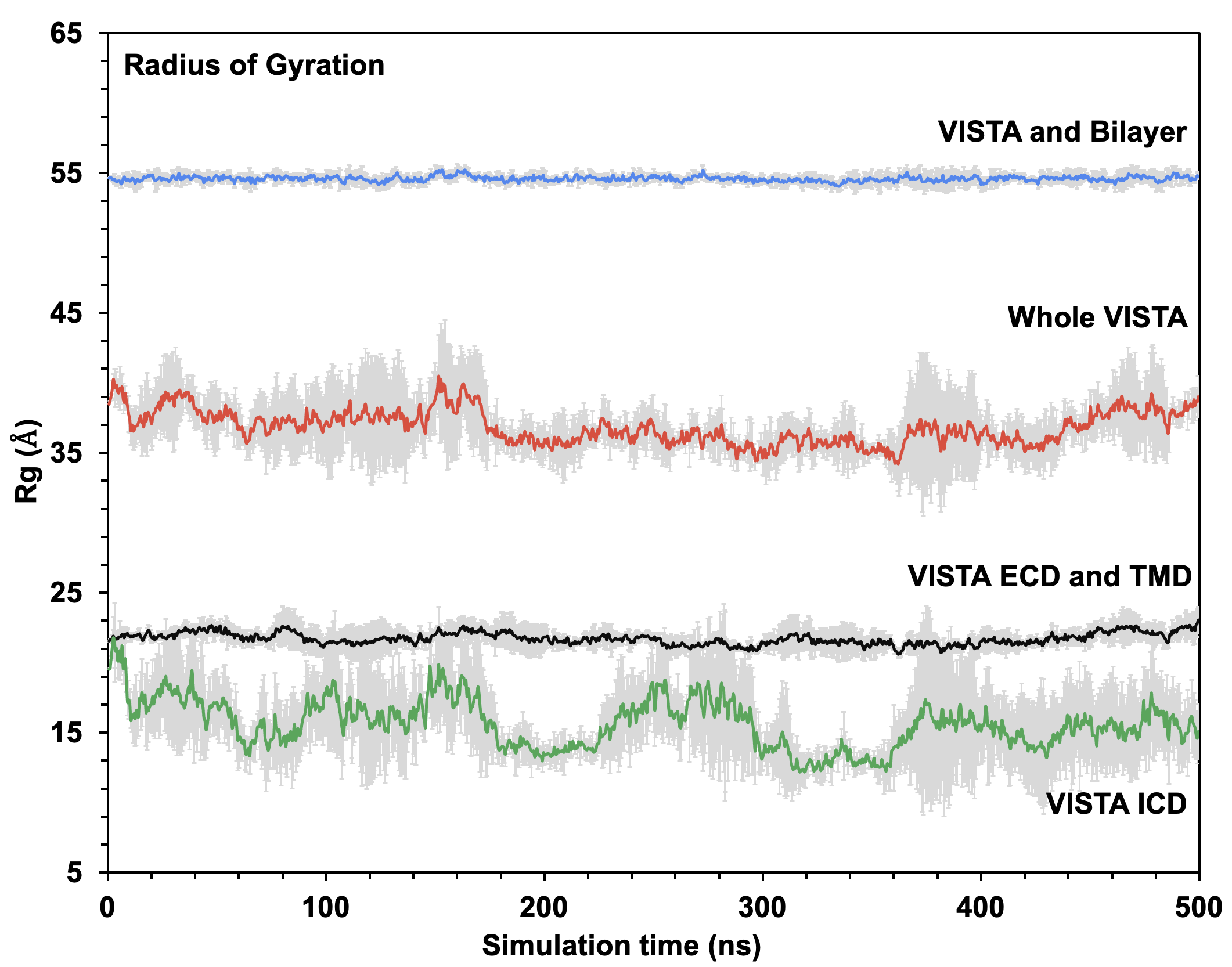

**Figure S4. The backbone RMSDs of different loops within the VISTA ECD, averaged over three MD replicates.** Most loops in the VISTA ECD remained relatively stable (less than 2.5 Å) during the MD simulations. The GH loop, however, exhibited highly dynamic behavior, with its backbone RMSD fluctuating up to 8 Å during the simulations. The ECD segment (residues 1 to 152) was first aligned before the RMSD of the individual loops were computed.

**Figure S5**

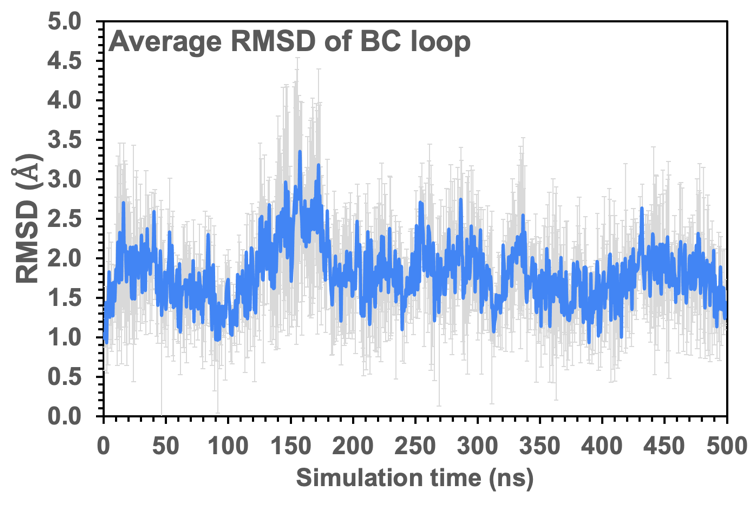

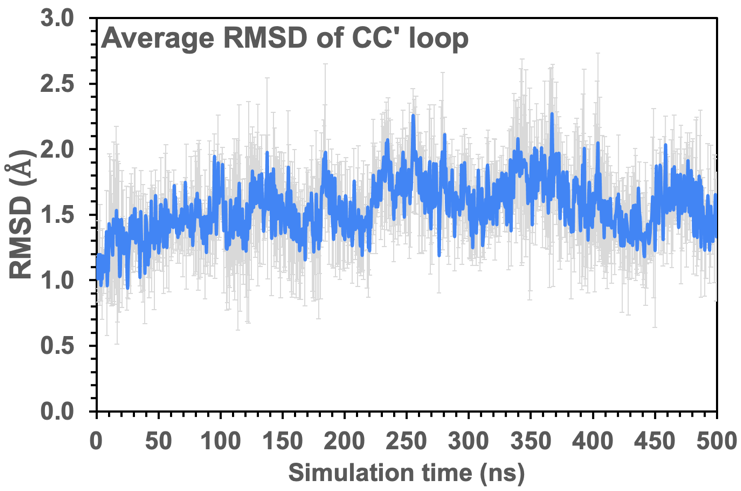

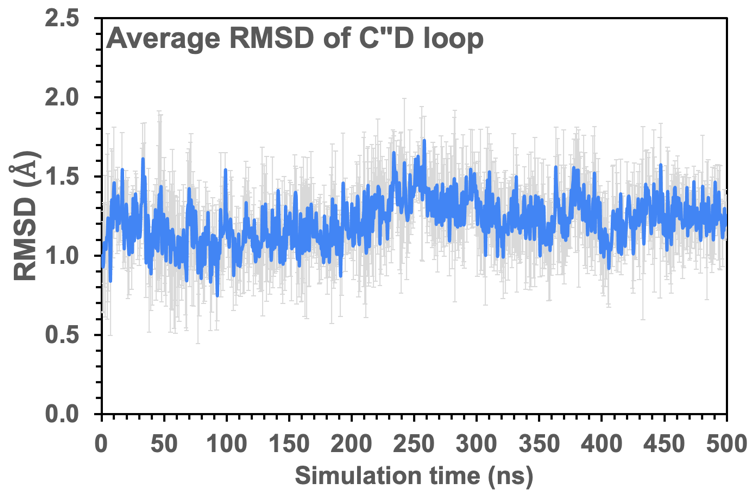

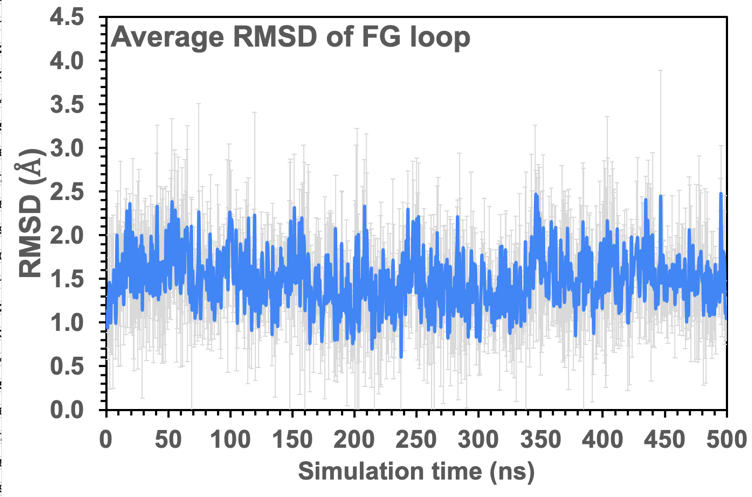

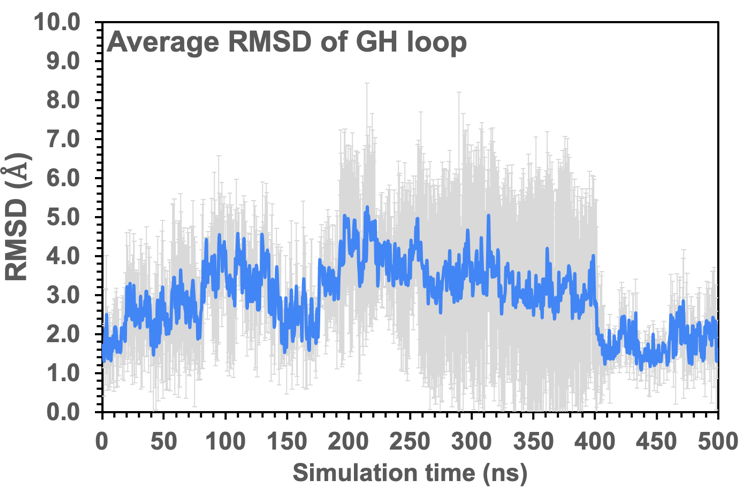

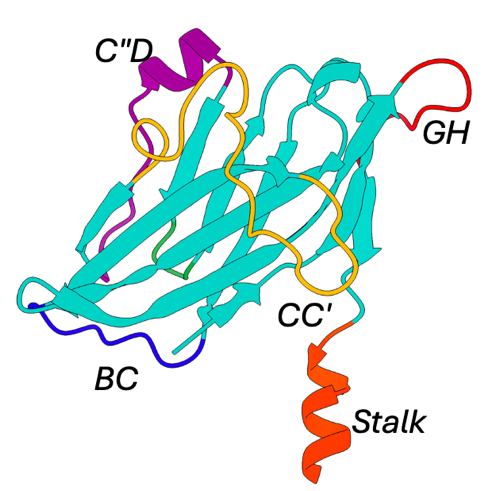

**Figure S5. Conformational stability of loops and inter-residue interactions in the VISTA ECD, averaged over three MD replicates.** The time evolution of the sveraged backbone RMSDs of the critical loops, BC, CC', C"D, FG, and GH, are shown. The CC', C"D, and FG loops remained highly stable, while the GH loop exhibited larger flexibility during the course of MD simulations.

**Figure S6:** Time evolution of inter-residue H-bond distances between residue pairs that stabilize the CC′ loop across all three MD simulation replicates.

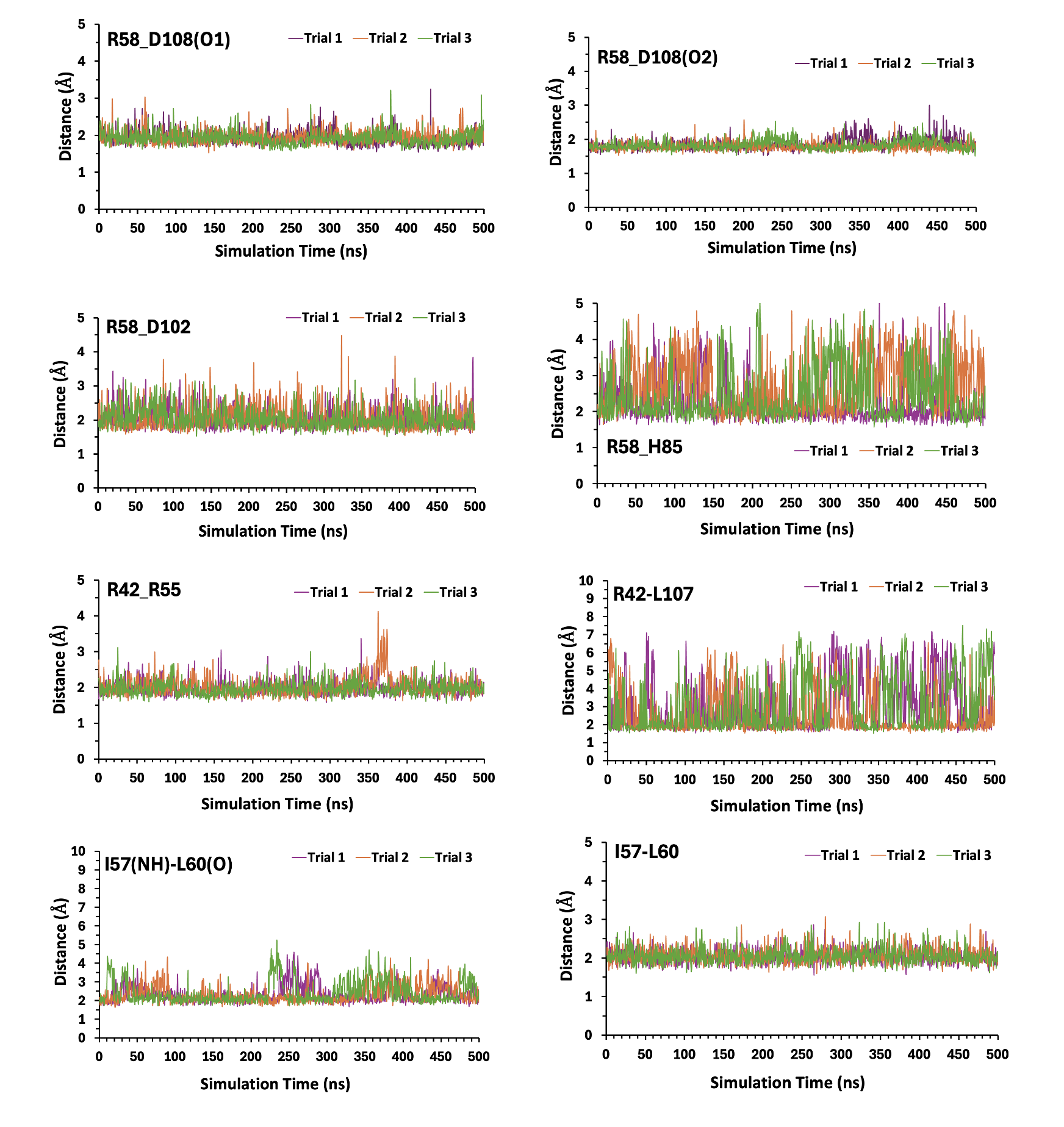

**Figure S6 (continued):** Time evolution of inter-residue H-bond distances between residue pairs that stabilize the CC′ loop across all three MD simulation replicates.

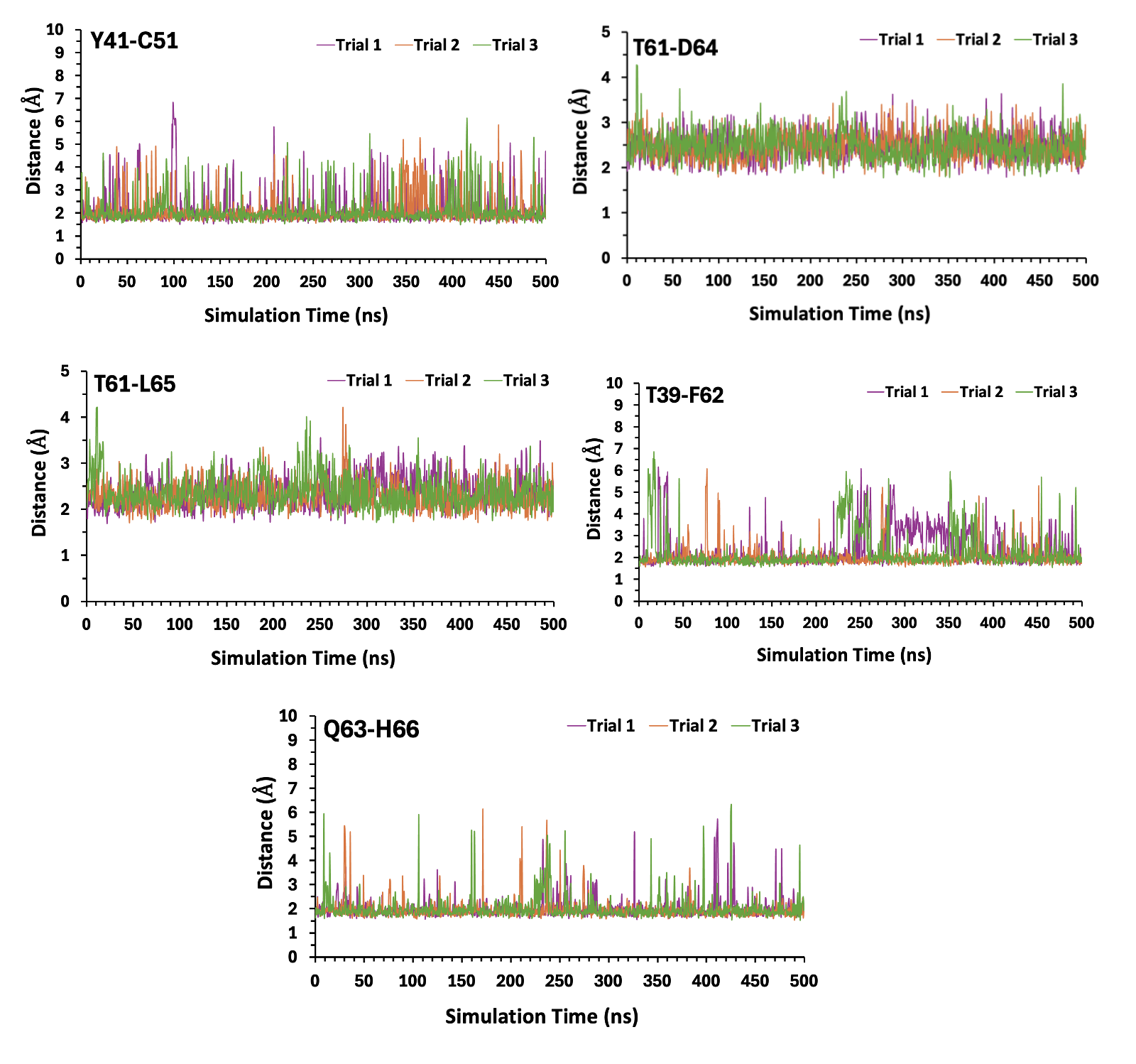

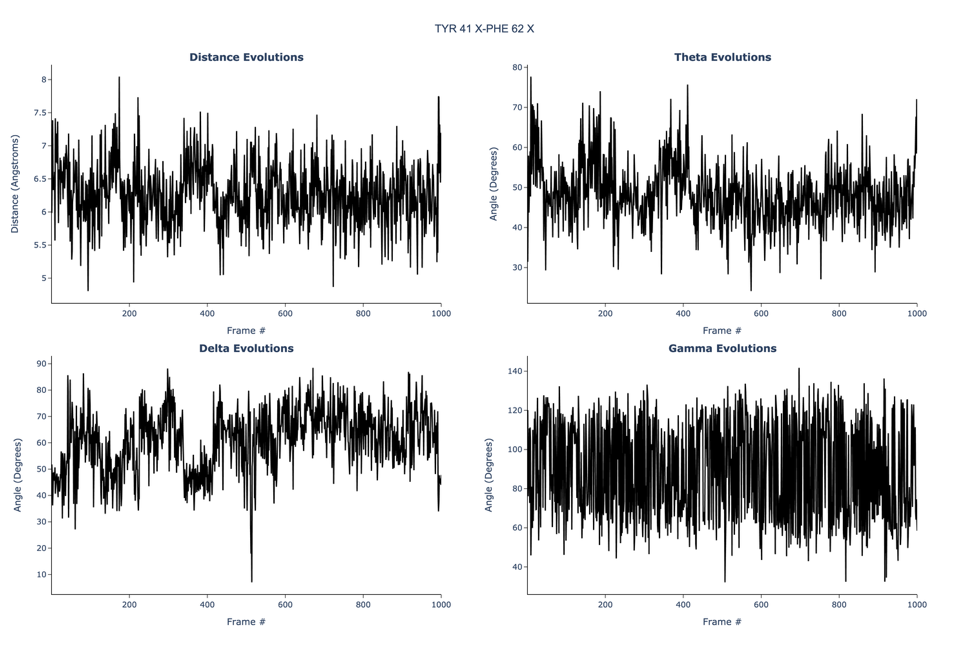

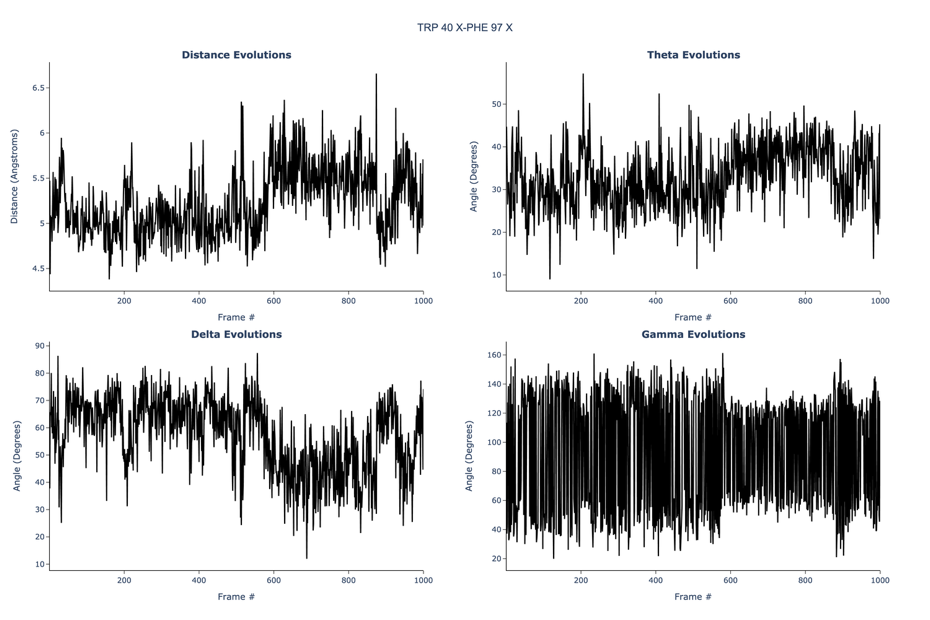

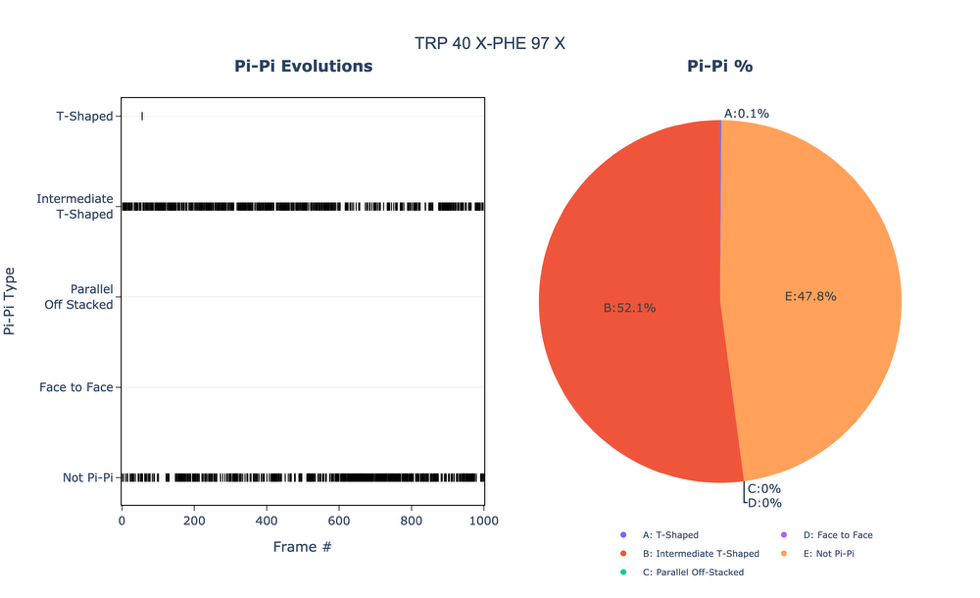

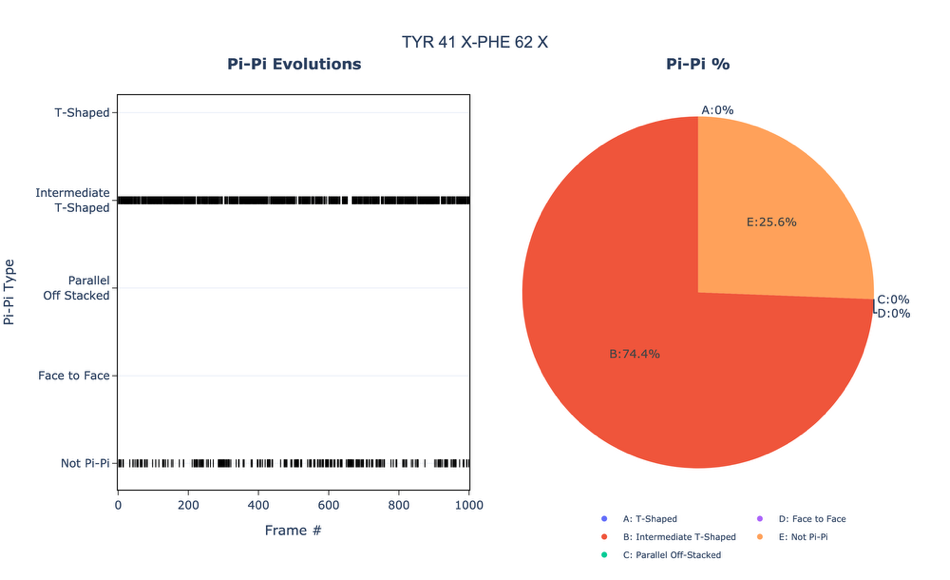
**Figure S7. Intermediate Stacking Interactions in the VISTA ECD, Assessed Across Three MD Replicates**. Two aromatic pairs in VISTA ECD were identified to exhibit intermediate-type of stacking interactions across the three MD replicates (trial 1 in A, trial 2 in B, and trail 3 in C). The evolution of the interaction types over the trajectory and a pie-chart for overall occupancy are shown in the left panels. These quantify the stability of the contact. The right panels show the time evolution of the key geometric parameters that define the stacking geometry: ring center distance (Å), and the three defining angles, Theta, Delta, and Gamma dihedrals, which together track the rings' sliding and rotational motions. X-axes shown as frame numbers, where 1 frame is equal to 0.5 ns.

(**A) MD trial=1**

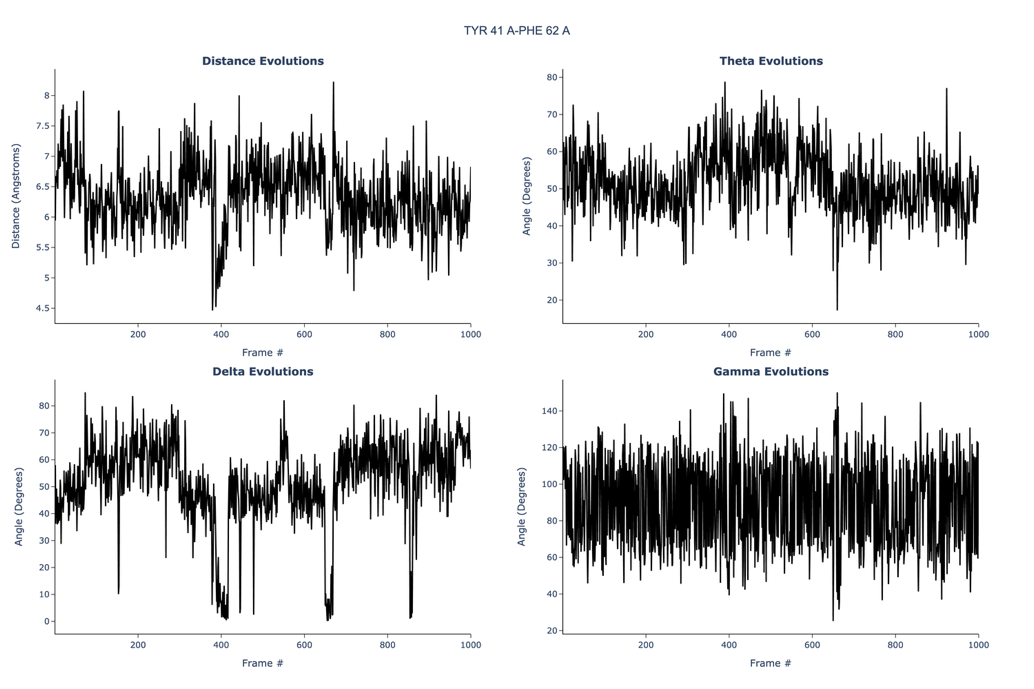

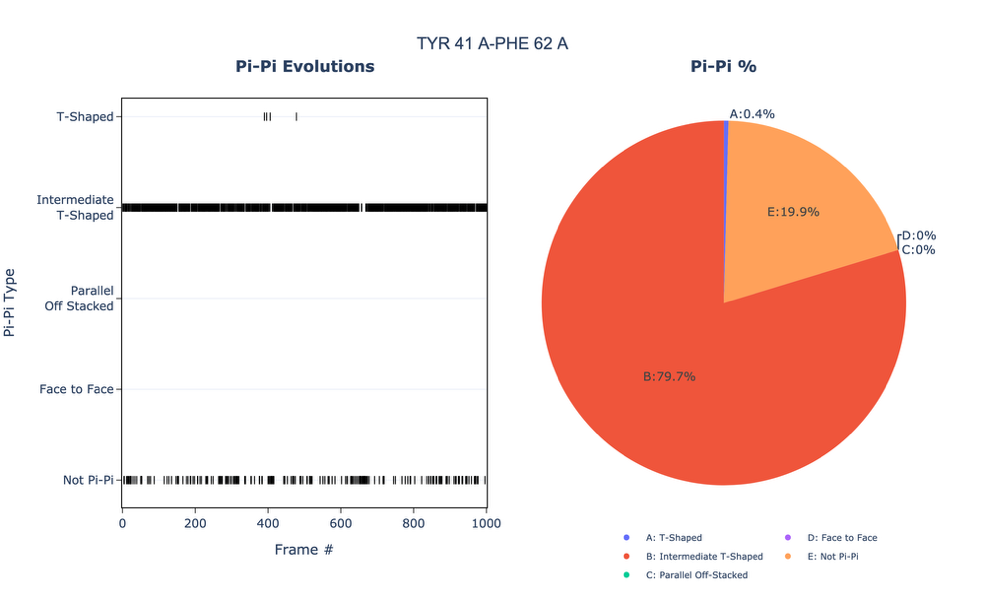
(**B) MD trial=2**

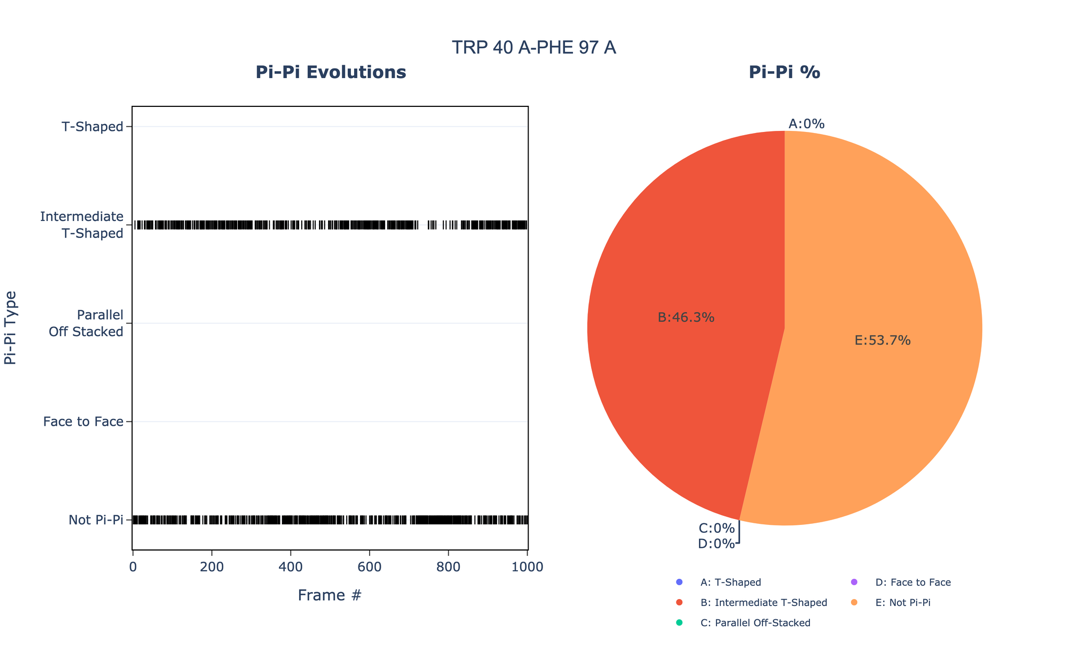

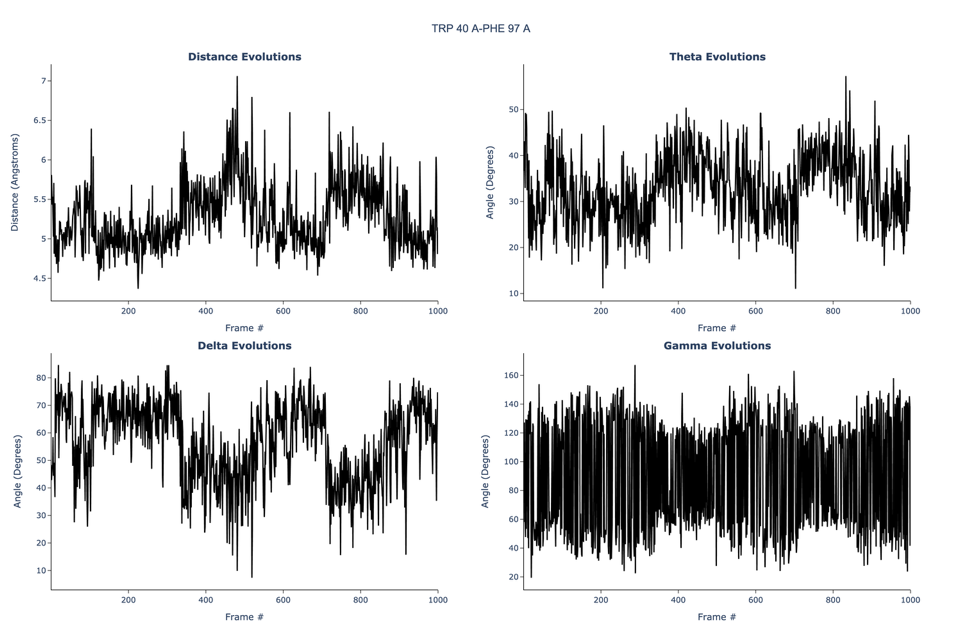

(**C) MD trial=3**

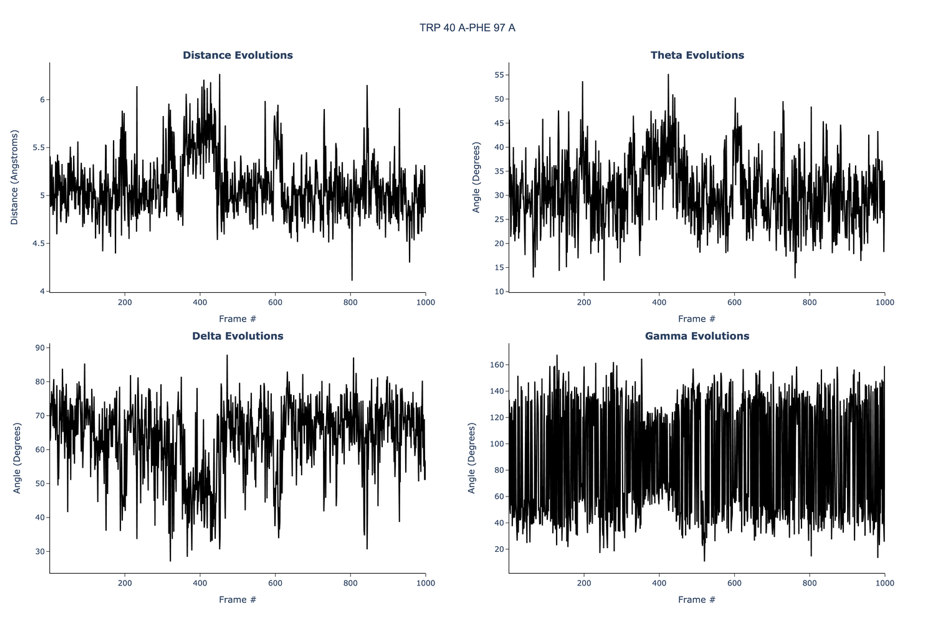

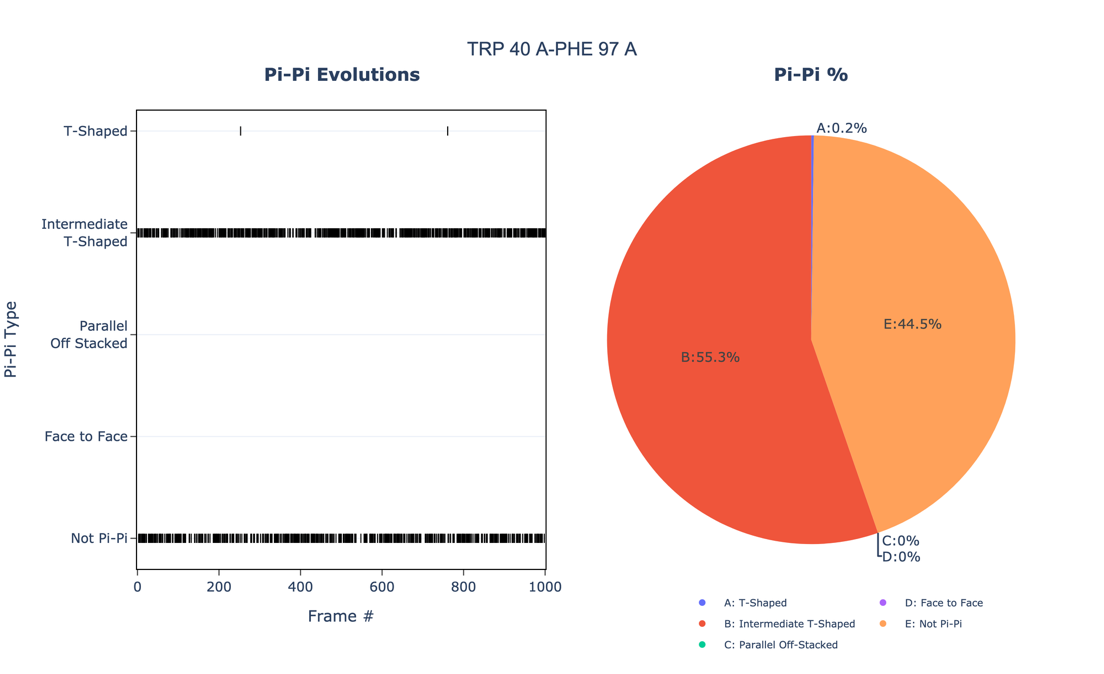

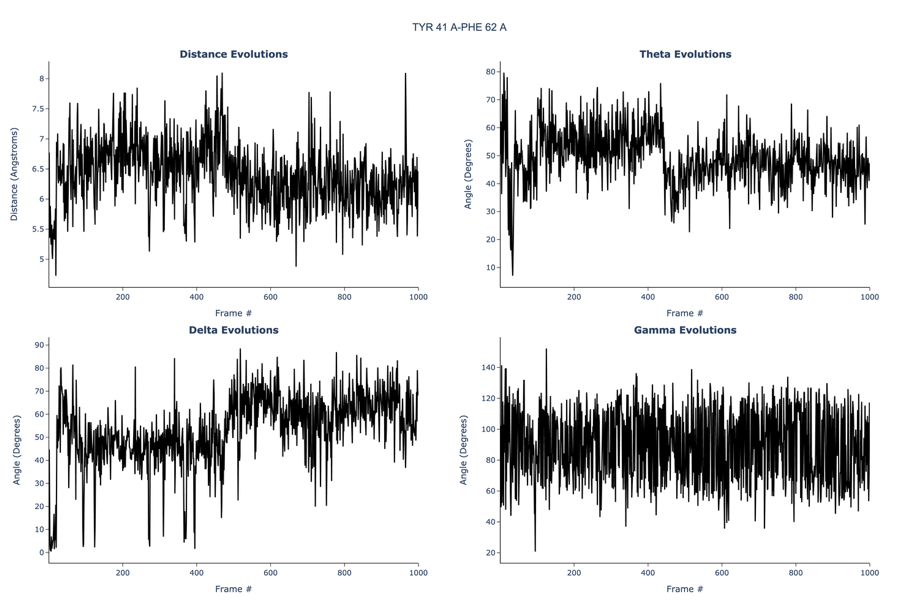

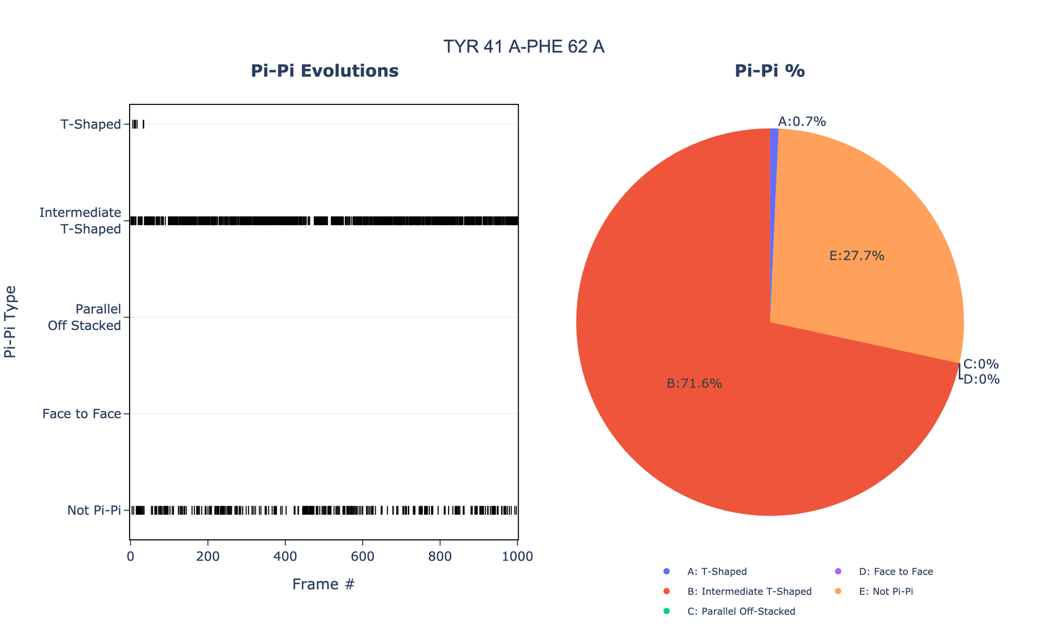

**Figure S8. A list of transmembrane (TM) sequences from the proteins within the CD28/B7 family of proteins. Proline residues in the TM sequences are highlighted in yellow.**

| Transmembrane sequences of CD28/B7 Family of proteins | | | |
| --- | --- | --- | --- |
| **Protein** | **Uniport ID** | | **Transmembrane Sequence** |
| VISTA_HUMAN | Q9H7M9 | | ALATGACIVGILCLPLILLLV |
| CD80_HUMAN | P33681 | | LLPSWAITLISVNGIFVICCL |
| CD86_HUMAN | P42081 | | WITAVLPTVIICVMVFCLILW |
| PD-L1_HUMAN | Q9NZQ7 | | THLVILGAILLCLGVALTFIF |
| PD-L2_HUMAN | Q9BQ51 | | WLLHIFIPFCIIAFIFIATVI |
| VSIG3_HUMAN | Q5DX21 | | LIAGAIGTGAVIIIFCIALIL |
| PSGL-1_HUMAN | Q14242 | | LLAILILALVATIFFVCTVVL |
| CD28_HUMAN | P10747 | FWVLVVVGGVLACYSLLVTVAFIIFWV | |
| CTLA-4 | P26410 | FLLWILAAVSSGLFFYSFLLT | |
| PD-1 | Q15116 | VGVVGGLLGSLVLLVWVLAVI | |
| ICOS | Q9Y6W8 | FWLPIGCAAFVVVCILGCILI | |
| TIGIT | Q495A1 | LLGAMAATLVVICTAVIVVVA | |
| TIM-3 | Q8TDQ0 | IYIGAGICAGLALALIFGALI | |

**Figure S9:**

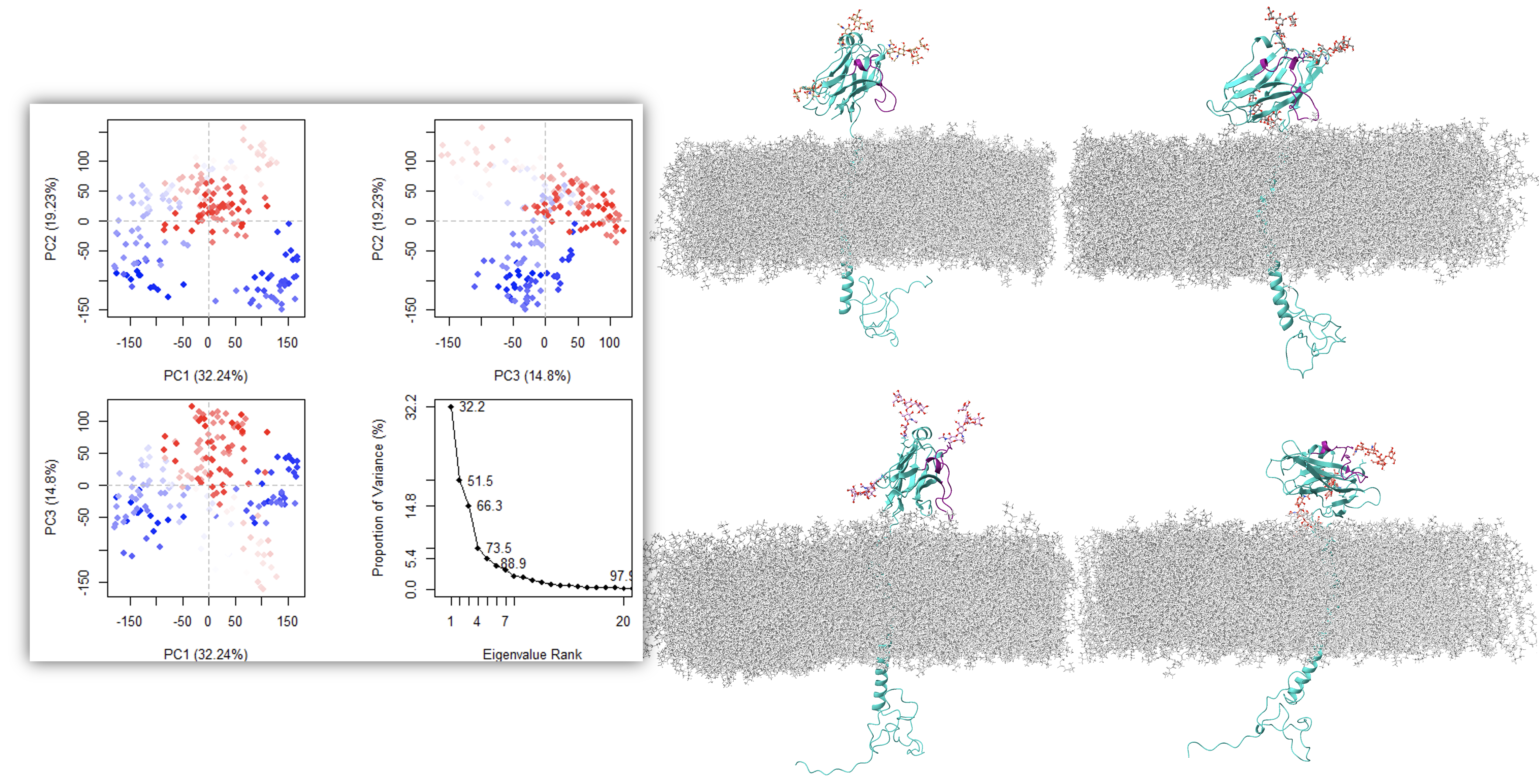

**Figure S9: PCA and conformational clustering to delineate essential dynamics of the glycosylated, and membrane-bound, full VISTA model essential dynamics during MD simulation.** The conformational landscape of VISTA is explored through the 2D projection of trajectories onto the first two principal components (PC1 and PC2). Distinct clusters in PCA plots are color-coded to represent the sampling of metastable states. Representative structures from each cluster are shown (right) with the CC' loop colored in purple and the remainder of the protein depicted in cyan ribbon. The glycans are shown as ball and stick representations. These snapshots reveal the two distinct conformations, up and down, that were identified in the glycan-free VISTA model. The magnitude of these essential motions is quantified via scree plots (left bottom), which illustrate the proportion of variance captured per eigenvalue rank. The dominance of the first few components underscores that the rotation of ECD constitutes the primary functional motion of the system.
